# Characterization of *Eh*PIG-K, a catalytic subunit of the GPI transamidase complex in *Entamoeba histolytica*

**DOI:** 10.64898/2026.11.24.768833

**Authors:** Simra Anees, Arnab Das, Zenab Kamali, Samudrala Gourinath

## Abstract

Glycosylphosphatidylinositol (GPI) anchors are essential for tethering surface proteins that mediate host-parasite interactions and contribute to the virulence of *Entamoeba histolytica*. The GPI transamidase (GPIT) complex, with PIG-K as its main catalytic enzyme, carries out the final step in GPI anchor addition. In this study, we examined the subcellular localization, enzymatic properties, and physiological importance of *E. histolytica* PIG-K (*Eh*PIG-K). Immunofluorescence microscopy showed that *Eh*PIG-K is primarily localized to a reticular network typical of the endoplasmic reticulum, around the nucleus and just beneath the plasma membrane, consistent with known sites of GPI anchor synthesis. The functional assay using a fluorescent peptide showed that *Eh*PIG-K has a peptidase activity with an optimum at pH 7.5. The presence of Ca^+2^ and Mg^+2^ ions greatly enhanced this activity whereas Mn^+2^ ion has a reducing effect. Ca^+2^ and Mg^+2^ ion binding release the inhibitory effect of the loop 199-223, adjacent to the active site residues, on the entrance to the catalytic nucleophilic core, whereas Mn^+2^ ion binding fixes the loop 199-223 in an inhibitory state. Suppressing the expression of *Eh*PIG-K significantly reduced cell growth, erythrophagocytosis, and cell survival compared to control and overexpressing cells. In addition, the level of Gal/GalNAc lectin, a major GPI- anchored cell-surface protein, was significantly reduced in *Eh*PIG-K-suppressed trophozoites. Overall, these findings show that *Eh*PIG-K is necessary for the cell-surface expression of GPI- anchored proteins and for key virulence-related functions, making *Eh*PIG-K a promising target for treating amoebiasis.

## Introduction

Amoebiasis is caused by *Entamoeba histolytica,* an intestinal protozoan parasite, and is a major health concern in many developing countries[1]. The parasite colonizes the large intestine, causes rupture of epithelial cells lining it, and reaches the liver and other major organs via the bloodstream. Its genome size consists of ∼24 Mbp [2]. Approximately 40,000 – 100,000 deaths per year have been reported due to this disease across the globe [1]. Many cell surface molecules of the *E. histolytica* trophozoite play important roles in regulating host cell adhesion, cytolysis, and pathogenesis [3]. One of these is a glycosylphosphatidylinositol (GPI) anchored protein, the Gal/GalNAc lectin, that abundantly coats the protozoan’s cell surface and is required for the binding of the trophozoites to host cell surface mucins [4]. Another important GPI anchored factor enriched in the glycocalyx layer is the proteophosphoglycan (PPG), possessing a divergent GPI anchor core [5].

Genome analysis of *E. histolytica* identified GPI biosynthesis genes including all core catalytic subunits of the enzymatic complex, compared with those of yeast and humans [6]. Studies using an antisense RNA approach against *Eh*PIG-L show that the amount of GPI- anchored proteins on the pathogen surface is reduced by 80-90%, which has a damaging effect on endocytosis, cell proliferation, fluid-phase pinocytosis, and adhesion to target cells [6]. A similar study using antisense RNA against *Eh*PIG-M, the first mannosyltransferase of the GPI biosynthetic pathway, also showed 85-90% reduction in GPIs at the cell surface [7]. Thus, enzymes of the GPI biosynthetic pathway are potentially interesting drug targets.

GPI biosynthesis is an important and ubiquitous pathway in all eukaryotes which produces a glycolipid anchor with a common conserved backbone consisting of ethanolamine-PO_4_-6Manα1-2Manα1-6Manα1-4GlcNα1-6*myo*-inositol-1-PO_4_-lipid [8]. Species-specific and tissue-specific modifications are built onto this core structure [9]. The GPI precursor is then transferred to the C-terminal end of proteins, transported to the Golgi apparatus and finally to the plasma membrane via the secretory pathway [9]. Proteins to be GPI anchored possess a C-terminal signal sequence (SS) of around 15-31 amino acids for GPI attachment with certain conserved features. The site at which the SS is cleaved and the protein is attached (ω residue), as well as two residues C-terminal to it (ω+1 and ω+2), are normally amino acids with small side chains. This is followed by a set of 5-10 polar residues and a hydrophobic domain comprising 15-20 hydrophobic amino acids [9].

GPI transamidase (GPIT) catalyses the cleavage of SS and transfer of the GPI precursors to proteins to generate GPI-anchored proteins [10]. In mammals and yeast, it is made up of 5 subunits, GPAA1(mammals)/Gaa1(Yeast), PIG-K/Gpi8, PIGT/Gpi16, PIGS/Gpi17, and PIGU/Gab1 [9]. *T. brucei* GPIT is also a multi-enzyme complex with two unique subunits, TTA1 and TTA2, and three other subunits, *Tb*GAA1, *Tb*GPI8, and *Tb*GPI16, which are homologous to their human counterparts (GAA1, PIG-K, and PIG-T, respectively) [11]. The published Cryo-EM structure of the human GPIT complex showed the arrangement of the five subunits, the interactions between them, molecular details of the active site, and that PIG-K functions as the catalytic component, which is important for the transamination reaction [12] [13]. In *T. brucei,* deletion of the GPI8/PIG-K subunit resulted in the absence of GPI-anchored proteins from the cell surface [14]. Strikingly, the initial bioinformatics and genome database searches for *Entamoeba histolytica* revealed homologs of four subunits (*Eh*PIG-K, *Eh*PIG-S, *Eh*PIG-T, and *Eh*PIG-U), but no obvious homolog of the GPAA1 subunit was found in the initial genomic survey [6]. A putative, uncharacterised protein corresponding to GPAA1 is, however, present in more recent pathway mapping (KEGG pathway annotations) and updated genome databases. This implies that the *E. histolytica* GPIT complex likely retains the fundamental architecture, including all five conserved subunits (*Eh*PIG-K, *Eh*PIG-S, *Eh*PIG- T, *Eh*PIG-U, and putative *Eh*GPAA1). Of these, *Eh*PIG-K is presumably the central catalytic subunit and contains the cysteine peptidase active site required for the SS cleavage. Sequence comparison shows that *Eh*PIG-K shares 36% identity with human (*Homo sapiens*) PIG-K, 38.17% with *Candida albicans* (Gpi8), and 34.63% with *Trypanosoma brucei*, showing a high degree of evolutionary conservation across species.

The molecular mechanism underlying this transamidation step remains uncharacterized in this parasite. Also, despite its clinical importance, the biochemical properties, intracellular localization, enzymatic activity, and essentiality of the catalytic subunit, *Eh*PIG-K, in *E. histolytica* remain completely unknown. The present study provides the first comprehensive biochemical, cellular, and functional characterization of *Eh*PIG-K. By combining in silico modelling, recombinant enzyme assays, inducible gene-suppression/overexpression, and confocal microscopy, this work reveals an essential role for *Eh*PIG-K in GPI-anchor processing, cell-surface presentation of major adhesins, cellular proliferation, and parasitic erythrophagocytosis.

## Material and methods

### *Entamoeba histolytica* culture, maintenance, and selection of the stable transfectants

*Entamoeba histolytica* (HM1:IMSS strain) and all transformed strains were cultured in TY1-S-33 medium supplemented with 15% adult bovine serum, 1X Diamond’s vitamin mix, 125 μl of 250 U/ml Benzyl Penicillin, and 0.25 mg/ml Streptomycin per 100 ml of medium. Cultures were maintained at 35.5°C under controlled conditions [15]. Stable cell lines expressing sense, antisense, and GFP-tagged *Eh*PIG-K were generated by electroporation. For electroporation, log-phase trophozoites were collected in PBS and washed with incomplete cytomix buffer (10 mM K_2_HPO_4_/KH_2_PO_4_, pH 7.6, 120 mM KCl, 0.15 mM CaCl_2_, 25 mM HEPES pH 7.4, 2 mM EGTA, and 5 mM MgCl_2_) [16]. Cells were resuspended in 0.8 ml of complete cytomix buffer, which consisted of incomplete cytomix supplemented with 4 mM ATP, 10 mM glutathione, and 150 μg plasmid DNA. The cell suspension was subjected to two electrical pulses at 3000 V/cm (1.2 kV) and 25 mF using a Bio-Rad electroporator. Following transfection, cells were initially cultured without selection. Selection and maintenance of stable transfectants began two days later, or upon reaching confluency, using increasing concentrations of G418 up to 10 μg/ml for constitutive expression constructs, or 10 μg/ml hygromycin B for tetracycline-inducible constructs. For experimental procedures, stable transfectants were grown to 50-70% confluency (or for up to 48 hours) and induced with 30 μg/ml G418 for GFP transformants or 30 μg/ml tetracycline for tetracycline-inducible transfectants for 36 hours.

### Cloning of various constructs used for *in vivo* and *in vitro* studies

For in vivo studies, the CAT gene from the pEhHYG-tet-O-CAT (TOC) shuttle vector was removed using KpnI and BamHI restriction enzymes. The *EhPIG-K* genes were then inserted in either the sense or antisense orientation to enable sense (S) or antisense (AS) RNA expression in *E*. *histolytica* trophozoites. For the GFP construct, the amplified full-length gene was cloned into the KpnI and EcoRV sites in the Neo-GFP vector, resulting in a C-terminus GFP tag.

The protein-coding regions of *EhPIG-K* (EHI_092280) were PCR-amplified from genomic DNA using primers containing NcoI and XhoI restriction sites for a C-terminus His tag. The respective DNA fragments were digested with the same restriction enzymes and further cloned into the T7 promoter-based pET-28a(+) expression vector for recombinant protein expression.

### Cell lysate preparation and western blotting

A total of 10^7^ logarithmic-phase cells were collected by centrifuging at 280 × g for 5 minutes at 4°C. The resulting pellet was washed with chilled PBS at pH 7.2. After washing, the cells were lysed using amoeba lysis buffer containing 50 mM Tris-HCl (pH 7.5), 150 mM NaCl, 1 mM PMSF, 0.5mM DTT, 1% Triton X-100, and 1X PIC. The lysate was kept on ice for 30 minutes, then centrifuged at 10,000 rpm for 30 minutes at 4°C to remove debris and unbroken cells. The protein concentration in the supernatant was measured using the BCA assay. Samples were then prepared in 1X SDS-PAGE loading dye. The measured amount of cell lysate was loaded onto a 12% SDS-PAGE gel with a prestained protein ladder (ThermoScientific), followed by transferring the protein from the gel via a semi-dry transfer system to polyvinylidene fluoride(PVDF) membrane using transferring buffer (25mM Tris, 192 mM Glycine, 20% (w/v) methanol, pH 8.3) at a constant 18V. The membrane was blocked with 5% BSA, followed by primary antibody incubation at 4^0^C anti-GFP (1:5000, rabbit-raised), anti-FLAG (1:3000, rabbit-raised), and anti-EhCabp1 (1:3000, mouse-raised). After primary antibody incubation, the blot was treated with secondary anti-rabbit 1:10000 or anti-mouse 1:5000 HRP-conjugated antibody (HRP, horseradish peroxidase) at room temperature for 1 hour, and then developed on a ChemiDoc using ECL substrate (Bio-Rad).

### Total RNA isolation and Real-time quantitative reverse transcriptase-PCR

The total RNA was extracted with the TRIzol reagent (Invitrogen). cDNA was prepared using the cDNA Synthesis kit (Thermo Scientific), and PCR was done with the primers listed in Table S2. The PCR reactions used Power SYBR Green PCR Master Mix (Applied Biosystems) on the 7500 Fast Real-Time PCR System (Applied Biosystems). The reaction steps were denaturation at 95 °C for 10 minutes, followed by 40 cycles at 95 °C for 15 seconds and 60 °C for 40 seconds. Melting curve analysis checked amplification specificity. Gene expression was measured using the ΔΔCT method.

### Protein Expression and Purification

The recombinant plasmid containing the *Eh*PIG-K gene was transformed into *E. coli* C41 (DE3) competent cells. For protein expression, a primary culture was grown by overnight incubation of a single transformed colony in LB medium. Subsequently, 1% of the primary culture was inoculated into Terrific Broth (Himedia) supplemented with 50 µg/mL kanamycin and allowed to grow at 37°C until the OD600 reached 0.8-1.0, then induced with 0.2 mM isopropyl β-D-1-thiogalactopyranoside (IPTG) and allowed to grow for 16 hours at 16°C. Cells were harvested by centrifugation at 8000 rpm for 10 minutes at 4°C. Harvested cells were resuspended and homogenized in lysis buffer containing 50 mM Tris-HCl (pH 7.5), 150 mM NaCl, 25 mM arginine, 0.1% Triton X-100, 0.1% CHAPS, 0.01% LMNG (Lauryl Maltose Neopentyl Glycol), 5 mM β-mercaptoethanol (β-ME), and 5% glycerol. Cell lysis was done by flash-freezing in liquid nitrogen and thawing in a 37°C water bath for up to five cycles. The lysate was then sonicated with 5–6 cycles (QSonica Ultrasonic Systems) on ice at 30% amplitude, with each cycle consisting of four to five 30-second pulses separated by 1-minute intervals. It was then centrifuged at 17,000 rpm for 30 minutes at 4°C. The supernatant was passed to a Ni-NTA column (GE Healthcare) pre-equilibrated with resuspension buffer (50 mM Tris-HCl pH 7.5, 150 mM NaCl, 5% glycerol, and 5 mM β-ME). After binding, the column was first washed with wash buffer 1 (50 mM Tris-HCl, pH 7.5, 150 mM NaCl, and 5 mM β-ME) and then wash buffer 2 (50 mM Tris-HCl, pH 7.5, 500 mM NaCl, 10 mM imidazole, and 5 mM β-ME). The bound protein was eluted with a buffer containing 50 mM Tris-HCl (pH 7.5), 150 mM NaCl, 25 mM arginine, 5 mM β-ME, and 300 mM imidazole. Protein homogeneity was evaluated by SDS-PAGE. Protein concentration was determined using the Bradford assay.

### Immunofluorescence staining

*E. histolytica* cells were collected by centrifugation, washed, and resuspended in TYI-33 incomplete medium [17]. The cells were allowed to adhere to acetone-cleaned coverslips in a 6-well plate for 30 minutes at 35.5 °C. After removing the medium, the cells were fixed with either 3% or 4% pre-warmed paraformaldehyde (PFA) for 30 minutes. The cells were then either permeabilised with 0.1% Triton X-100 in PBS for 10 minutes or washed with PBS and quenched for 30 minutes in PBS containing 50 mM NH_4_Cl. Next, the cells were blocked with 1% BSA in PBS and incubated with the primary antibody at 37 °C for 2 hours. After washing with PBS and 1% BSA, the cells were incubated with Alexa Fluor anti-mouse or anti-rabbit secondary antibody for 1 hour at 37 °C. The primary antibody dilutions were: anti-FLAG at 1:300, anti-*Eh*Rab7a at 1:200, anti-*Eh*CRT at 1:200, and anti-*Eh*Gal/GalNAc lectin at 1:150. To stain the ER or lysosomal compartments, cells were incubated with ER-Tracker^TM^ Red or LysoTracker^TM^ Red DND-99 (Cat. L-7528, Life Technologies, USA), respectively, for 2 hours at 35.5 °C. Cells were washed with PBS and mounted on glass slides using DABCO (1,4-diazbicyclo(2,2,2)octane, Sigma). The edges of the coverslip were sealed with nail polish to prevent drying. Confocal images were taken using a Nikon Real Time Laser Scanning Confocal Microscope, and raw images were processed with NIS-Elements 3.20 (Nikon).

### Live cell imaging

About 5×10^5^ GFP-*Eh*PIG-K induced cells were washed with PBS, incubated in incomplete medium, and placed on a 35 mm glass-bottom dish. Unstained RBCs were then added, and the dish was kept at 37°C on a temperature-controlled stage of a spinning-disk confocal microscope (Nikon A1R; Optics: Plan Apo VC606 oil DIC N2; Camera: Nikon A1; NA 1.4; RI 1.515). We collected differential interference contrast (DIC) and fluorescent time-lapse images of a moving amoeba. The raw images were analysed using NIS-Elements 3.20 software.

### Growth kinetics of *E. histolytica* trophozoites

The growth kinetics was performed by counted the cells using using a hemocytometer and then incubating an equal no. of cells (3*10^4^) for each construct in 7ml culture tubes. The cells were then allowed to grow at 35.5°C for 12, 24, 36, 48, and 72 hours. For each time point, cells were harvested by first chilling the tube on ice, then washing the cells with 1x PBS (pH 7.2) by centrifugation at 1500g for 5 min at 4^0^C, then resuspending the cells in PBS and counting them with a hemocytometer. The experiment was performed in triplicate, and the results are presented as mean ± standard error (SE).

### Erythrophagocytosis assay

The RBCs and the amoebic trophozoites for each construct were first washed using PBS at pH 7.2 and then resuspended in TYI-33 medium. The same number of cells (10^5^) was used, and the cells were incubated with RBCs at a 1:100 ratio at 37 °C for various time points. After incubation, the amoebae and erythrocytes were collected by centrifugation to remove unengulfed RBCs and then cold distilled water was added to the pellet, and the mixture was centrifuged at 1000 × g for 2 minutes. This washing step was carried out twice. The pellet thus obtained was resuspended in 1 mL of formic acid to lyse the amoebic trophozoites and release the RBCs. Absorbance was measured at 400 nm with a formic acid blank using a spectrophotometer. The experiment was performed in triplicate, and the results are given as the mean ± standard error (SE).

### Activity assay

The enzymatic activity of *Eh*PIG-K was measured in a reaction containing purified *Eh*PIG-K protein (2 mg/mL), 100 mM buffer (pH 5.0–8.5), and 10 μM peptide substrate. The reactions were incubated at 37 °C for 10 h. For the metal-ion-dependent assay, the respective divalent cation was added to a final concentration of 5 mM. After incubation, the reaction was centrifuged at 8,000 × g for 15 min. The supernatant thus obtained was collected, diluted as necessary with autoclaved distilled water, and then measured with a spectrofluorometer (Model RF-5301PC, Shimadzu). The fluorescence excitation at 360 nm and fluorescence emission from the released AMC were measured between 380 and 550 nm with slit widths of 3 nm for both excitation and emission. The experiment was performed in triplicate, and the results are presented as mean ± standard error (SE).

### Model generation for the *Eh*PIG-K

We used the AlphaFold Server3 [18] to generate PDBs for all four states of *Eh*PIG-K, namely Apo, Ca^+2^, Mg^+2^, and Mn^+2^-bound. We inspected the initial model and truncated it at the terminals to remove noise generated by the flexible disorder regions in the molecular dynamics simulation runs, owing to their highly fluctuating behaviour, and prepared the construct Apo *Eh*PIG-K_(20 - 279)_ and its 3 ion-bound states using PyMOL 3.1.1 [Schrödinger, LLC. *The PyMOL Molecular Graphics System, Version 3.1.1]*. We aligned all four models against the CryoEM reference structure of PIG-K in Human GPIT (PDB ID: 7WLD). They compared well with the reference structure and were used as the initial frames for the dynamic simulations.

### Molecular dynamics simulations

Total of 5 dynamic simulation runs for each of the four states of the *Eh*PIG-K were performed using the GROMACS 2023.2 package [19][20]. In the first step, the PDB structures were converted into GROMACS format using the pdb2gmx module. Then we used the CHARMM36 force field to generate protein topologies, including parameters for the bound ions, namely Ca^+2^, Mg^+2^, and Mn^+2^. Then, in the next step, we solvated the system using a cubic solvation box with periodic boundary conditions, setting the protein-boundary distance to 1.2 nm, and used the TIP3P water model for solvation. Thereafter, the systems were neutralised by addition of Na^+^ and Cl^-^ ions. After that, we minimised the energy of the systems using the steepest-descent algorithm (with a maximum of 50000 steps and an initial step size of 0.01nm) with a Verlet-cutoff scheme, until the maximum force fell below 1000.0 kJ/mol/nm. Energy minimized systems were then processed through NVT and NPT method, for temperature coupling we used modified Barendsen thermostat [21], set to 1.0 ps, maintained the temperature at 300 K, and for the isotropic press coupling used Parrinello-Rahman barostat (compressibility of 4.5 * 10^-5^ bar^-1^) at 2.0 time step. Hydrogen-bonded interactions were constrained using the LINCS algorithm [22]. The Particle Mesh Ewald system was used to treat the long-range electrostatic (Coulomb) interactions, while 1.0 nm cutoffs were set for short-range electrostatic and Van der Waals interactions [23]. For each run, the gen_seed of the NVT method was set to −1. Finally, a 100 ns simulation was run for each of the 5 replicas of each of the four states. Data were analysed using GROMACS; structures were visualised using Pymol 3.1.1. Root mean square deviation (RMSD), root mean square fluctuation (RMSF), radius of gyration (RG), solvent accessible surface area (SASA), energy and distance calculations were done using RMS, RMSF, RG, SASA, energy and distance modules provided by GROMACS, and plots were prepared using Matplotlib [24].

### Cartesian principal component analysis

For the whole protein Cartesian PCA, we used the 260 backbone Cα-atoms for the least-squares fitting and generation of the eigenvectors, and for the loop-specific Cartesian PCA. We performed least-squares fitting on carefully chosen core Cα-atoms that exhibited the least movement in the whole protein PCA, and for analysis and eigenvector generation, we used loop-specific Cα-atoms. For covariance analysis we used GROMACS covar module. To generate the models for the movement of the loops (Fig 10B, 10C, 11C, 11D and 11E) we used pymol trace atoms module to connect the Cα-atoms [25]. For the Gibbs free energy analysis, we used GROMACS sham module, and used python scripts for the for generation of the free energy landscape plots. First 40ns of each run were discarded to choose post equilibration frames, kept the criteria of choosing the bins shared by 15% by at least 3runs for each of the state, with no single run exceeding 50% in contribution. Used GROMOS cluster algorithm for the clustering. For analysing the trajectories we used MD analysis module [26][27] and for generations of the projections and time-resolved trajectories (Supplementary figure 8, 9, 11, 12, 13) used NumPy module of Python [28].

### Peptide docking

Used HADDOCK 2.4 for the docking of the peptide on 100 ns mds resolved structure of each state [29][30].

## Results

### 1. Expression and ER-Specific Localization of *Eh*PIG-K in *E. histolytica*

*Eh*PIG-K was FLAG-tagged at the C-terminus of the pEhHYG-tet-O-CAT (TOC) shuttle vector and stably transfected into wild type HM1:IMSS strain of *Entamoeba histolytica*. The expression of fused protein was confirmed through western blot of total cell lysates from transfected *E. histolytica* trophozoites that showed a clear band at approximately 35 kDa with an anti-FLAG antibody, which matches the expected size of the fusion protein (Figure 1A). Next, we used confocal fluorescence microscopy to determine the localisation of *Eh*PIG-K. The method used to fix the cells influenced the protein’s structure [31] [32]. When we fixed the cells with 3% paraformaldehyde (PFA), *Eh*PIG-K showed a clear punctate vesicular pattern that strongly overlapped with the ER tracker (Figure 1B). Quantitative pixel-intensity analysis yielded a high Pearson correlation coefficient (r ≈ 0.8), confirming that *Eh*PIG-K is primarily localised to the endoplasmic reticulum (ER). When we fixed the cells with 4% PFA, *Eh*PIG-K predominantly localizes to a reticular network characteristic of the ER, extending throughout the perinuclear region and beneath the plasma membrane, consistent with sites of GPI anchor biosynthesis (Figure 1D). To assess whether the signal was specific to organelles, we examined the overlap between *Eh*PIG-K and the DAPI-stained nucleus. The Pearson correlation coefficient was very low (r ≈ 0.1, Figure 1E), which rules out accumulation in the nucleus and shows that *Eh*PIG-K is anchored to the endoplasmic reticulum network and the nuclear envelope membrane that connects to it in *E. histolytica*.

**Figure 1:**
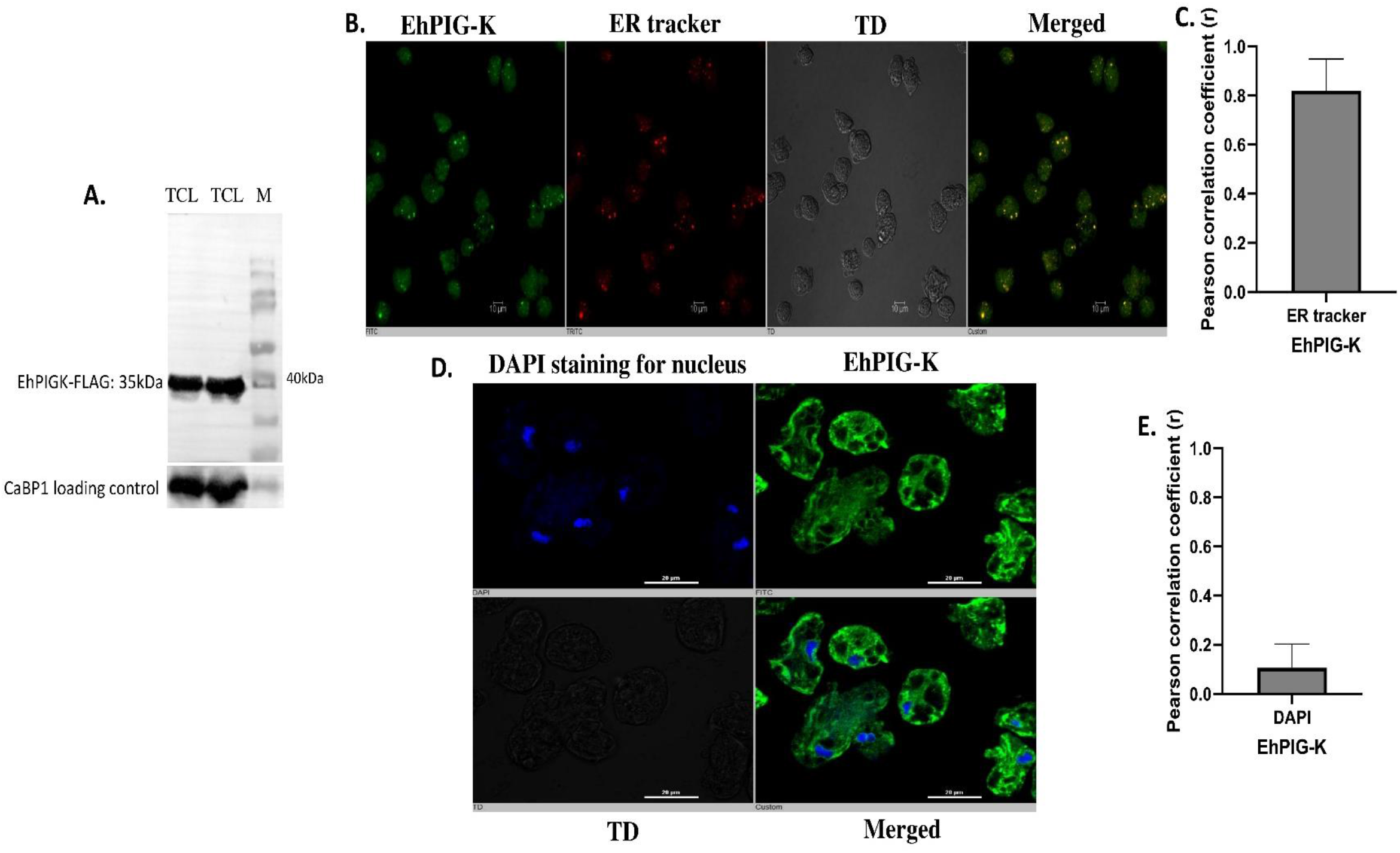
The expression, subcellular localisation and organelle specificity of *Eh*PIG-K in the trophozoites of *E. histolytica.* **(A)** Detection by western blotting of the FLAG-tagged *Eh*PIG-K in the total cell lysates of *Eh*PIG-K FLAG transfected cell line induced with 30 μg/ml tetracycline (TCL loaded as lane 1: 75 μg and lane 2: 100 μg) at about 35 kDa, taking CaBP1 as the loading control. **(B)** Confocal micrographs illustrating the co-localisation of *Eh*PIG-K (green) with the ER tracker (red) after fixation in 3% PFA. **(C)** The Pearson correlation coefficient (r) used to measure the overlap between the *Eh*PIG-K signal and the ER tracker (r approximately 0.8). **(D)** The subcellular distribution of *Eh*PIG-K (green) in relation to the nuclei stained with DAPI (blue) following fixation in 4% PFA, showing *a reticular network characteristic of the endoplasmic reticulum, extending throughout the perinuclear region and beneath the plasma membrane,* the nuclear envelope, and in the regions sub-plasma membrane. **(E)** The quantification of the co-localisation between *Eh*PIG-K and DAPI. Scale bars are shown in each panel. Quantification analysis was done from 30 cells for each respective panel. The bars indicate the mean ± SD.

### 2. *Eh*PIG-K Resides Intracellularly within the ER Network without Cell-Surface Exposure

To better understand where *Eh*PIG-K is located inside the cell and whether it appears at the cell surface or is associated with it, we used confocal immunofluorescence assays to compare *E. histolytica* trophozoites that were either left intact or made permeable with Triton X-100 with known ER proteins. We first checked whether *Eh*PIG-K is localised to the same region as Calreticulin (*Eh*CRT), an ER protein involved in the folding of glycosylated proteins, and it has also been reported to be present on the plasma membrane and to participate in phagocytosis in *E. histolytica* [33], which marks the endoplasmic reticulum in *E. histolytica*. In cells that were not made permeable, we did not see *Eh*PIG-K on the surface, but *Eh*CRT was present in the plasma membrane of the phagocytosed cells (top panel of Figure 2A). When we made the cells permeable with detergent, the antibodies could enter, and we observed a clear overlap between *Eh*PIG-K (green) and *Eh*CRT (red), forming a distinct reticular network characteristic of the endoplasmic reticulum (bottom panel of Figure 2A). Analysing the intensity of the signals in permeabilised cells gave a Pearson correlation coefficient of about 0.7 (Figure 2B), which means there is significant co-localization with the ER. Next, we checked whether *Eh*PIG-K is co-localised with the cell-surface lectin *Eh*Gal/GalNAc. In cells that were not permeabilised, *Eh*Gal/GalNAc marked the plasma membrane surface, but we did not detect *Eh*PIG-K on the cell surface (top panel of Figure 2C). After permeabilising the cells, we observed that *Eh*PIG-K (blue) and *Eh*Gal/GalNAc (green) overlapped within the ER and GPI- anchor assembly site (bottom panel of Figure 2C). The Pearson coefficient was about 0.7 (Figure 2D). Overall, these results show that *Eh*PIG-K is an intracellular protein anchored in the ER reticular network and is not present on the cell surface.

**Figure 2:**
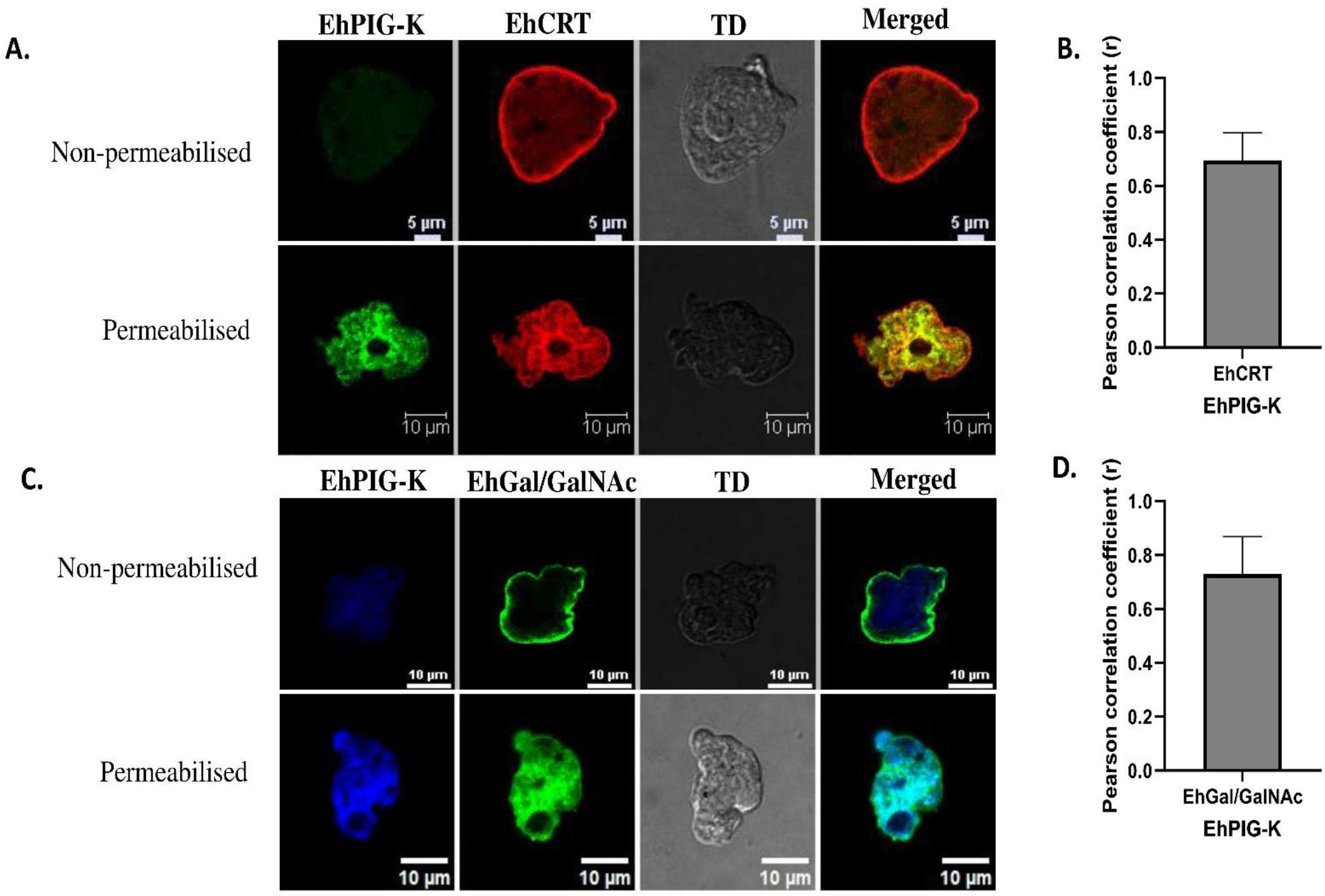
Co-localization of *Eh*PIG-K with ER marker *Eh*CRT and surface marker *Eh*Gal/GalNAc lectin. **(A)** Immunofluorescence assay of non-permeabilised and permeabilised *E. histolytica* trophozoites stained for *Eh*PIG-K (green) and the known ER marker *Eh*CRT (red). In non-permeabilised cells, there was no surface staining for *Eh*PIG-K but *Eh*CRT on the surface. After permeabilisation, clear intracellular co-localization with EhCRT was seen. **(B)** Co-localization between *Eh*PIG-K and *Eh*CRT in permeabilised cells was measured, with a Pearson correlation coefficient of about ≈ 0.7. **(C)** Confocal images of non-permeabilised and permeabilised trophozoites stained for *Eh*PIG-K (blue) and the surface lectin *Eh*Gal/GalNAc (green). In non-permeabilised cells, *Eh*Gal/GalNAc was found on the surface, but *Eh*PIG-K was not detected on the surface. In permeabilised trophozoites, the two signals overlapped inside the cell. **(D)** The Pearson correlation coefficient ≈ 0.7 indicates a spatial correlation between *Eh*PIG-K and *Eh*Gal/GalNAc in permeabilised trophozoites. Scale bars are shown in each panel. Quantification analysis was done from 30 cells for each respective panel. The bars indicate the mean ± SD.

### 3. *Eh*PIG-K Does Not Colocalize with Endosomal Marker *Eh*Rab7a and Lysosomal Marker (LysoTracker) in *Entamoeba histolytica*

*Entamoeba histolytica* displays a fluid and interconnected continuum that includes the endoplasmic reticulum (ER), Golgi-like structures, and dynamic vesicular networks. We used confocal microscopy to investigate the potential involvement of *Eh*PIG-K in the endolysosomal pathway. We examined the localization of *Eh*PIG-K and the late endosomal marker *Eh*Rab7a within the cell [34]. Immunofluorescence revealed that *Eh*PIG-K (green) and *Eh*Rab7a (red) are localised to distinct regions and do not colocalize (Figure 3A). The Pearson’s correlation coefficient was r ≈ 0.03, which confirms there is no significant colocalization between *Eh*PIG-K and late endosomal compartments (Figure 3B). Next, to determine whether *Eh*PIG-K is present in acidic lysosomal vesicles during phagocytosis, which represents a fundamental mechanism underlying tissue invasion and pathogenesis. We performed a time-course erythrophagocytosis assay using LysoTracker, a membrane diffusible probe that stains the acidic organelles [35]. Trophozoites expressing *Eh*PIG-K (green) and stained with LysoTracker (red) were mixed with red blood cells and analyzed at 0, 10, 20, and 30 minutes (Figure 3C). At all the time points, *Eh*PIG-K stayed separate from the LysoTracker-labelled lysosomes and phagolysosomal vacuoles that forms by the fusion of maturating late phagsomes to lysosomes for the degradation of ingested host cells, or red blood cells [36]. The Pearson’s correlation coefficient was about r ≈ 0.04, showing there was no overlap (Figure 3D). Overall, these results show that *Eh*PIG-K is not found in late endosomes or acidic lysosomal compartments, either when the cells are at rest or during erythrophagocytosis.

**Figure 3:**
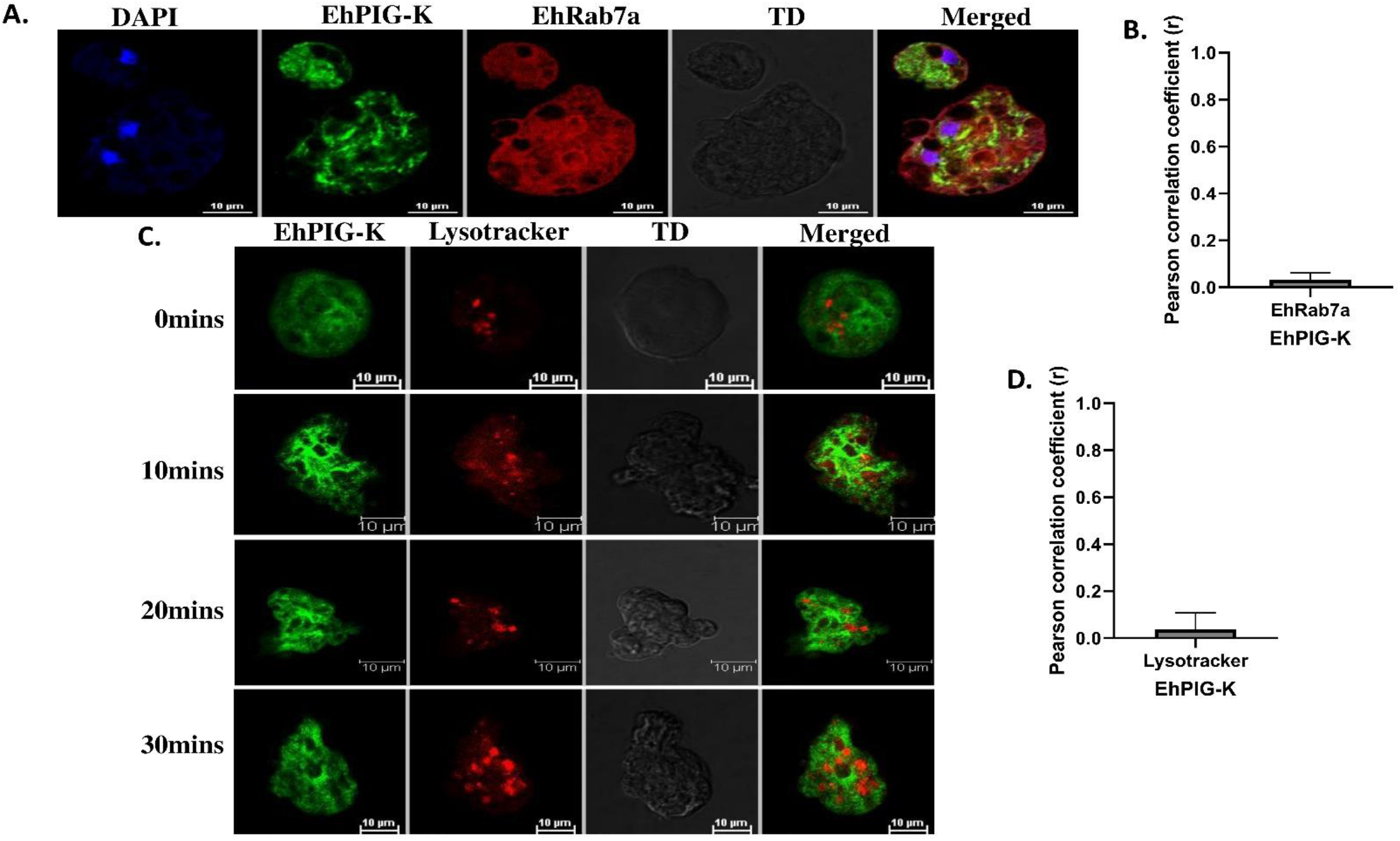
The subcellular localisation of *Eh*PIG-K and the extent of its overlap with markers of specific organelles. In **(A)**, immunofluorescence images of *E. histolytica* trophozoites are shown with *Eh*PIG-K stained in green and *Eh*Rab7a, a late endosomal GTPase, stained in red; the nuclei are labelled with DAPI (blue) and the brightfield images were obtained with the transmitted detector (TD). In **(B)**, a quantitative assessment of the colocalization between *Eh*PIG-K and *Eh*Rab7a gives a Pearson’s correlation coefficient (r ≈ 0.03), which indicates that there is no significant overlap. **(C)** Displays a series of time-course fluorescence images of trophozoites expressing *Eh*PIG-K (green) that have been incubated with LysoTracker (red) and erythrophagocytosis at 0, 10, 20, and 30 minutes. **(D)** The Pearson’s correlation coefficient (r ≈ 0.04) shows that there is no colocalization between *Eh*PIG-K and LysoTracker at any of the time points. Quantification analysis was done from 10 cells for each respective panel. The scale bars are 10 μm and the values are given as the mean ± SD from representative cells.

### 4. *Eh*PIG-K gene is necessary for cell proliferation, cell survival, and phagocytosis

To examine the functional role and biological importance of *Eh*PIG-K in *Entamoeba histolytica*, gene expression constructs were prepared for both the sense (leading to overexpression) and antisense (resulting in gene suppression) directions using a tetracycline-inducible Tet-O-CAT vector system (Figure 4A). qRT-PCR (Quantitative Real-Time Reverse Transcription PCR) analysis was used to verify that expression had been successfully modified: treatment with tetracycline caused a substantial increase (7-8 fold) in the relative mRNA levels in the overexpression cell line (sense), while the Antisense line showed a significant decrease in transcript abundance when compared with the control strains (Figure 4B). To further validate the biological effect of modulating *Eh*PIG-K on cellular growth, the growth kinetics of cell proliferation were observed over a period of 72 hours (Figure 4C). For this study, fresh growth media was inoculated with an equal number of cells and induced with tetracycline. At the designated time points, the cells were collected, re-suspended in PBS, and counted on a hemocytometer. Compared to the uninduced controls (SENSE UI, A/S UI) and to the vector control strains (TOC), the tetracycline-induced overexpression of *Eh*PIG-K (SENSE IN) led to a significant increase in cell proliferation. On the other hand, suppression of *Eh*PIG-K (A/S IN) greatly reduced cell growth and significantly inhibited trophozoite proliferation by 90%. We found that gene expression of *Eh*PIG-K is crucial for cell survivability and proliferation. Functional assays were also carried out to assess the impact of *Eh*PIG-K expression on the ability of the cells to carry out erythrophagocytosis, engulfment of RBCs, and to illustrate the role of *Eh*PIG-K in tissue invasion phenomenon (Figure 4D). The trophozoites that overexpressed *Eh*PIG-K (SENSE IN) showed a marked increase in their uptake capacity over time when compared with the control group. In contrast, when *Eh*PIG-K was suppressed (A/S IN), there was a substantial decrease in phagocytic activity by 90%, which shows that the level of *Eh*PIG-K expression is directly related to phagocytic efficiency in *E. histolytica*. This further supports the growth study data that *Eh*PIG-K is crucial and necessary for cell proliferation, motility, and phagocytosis.

**Figure 4:**
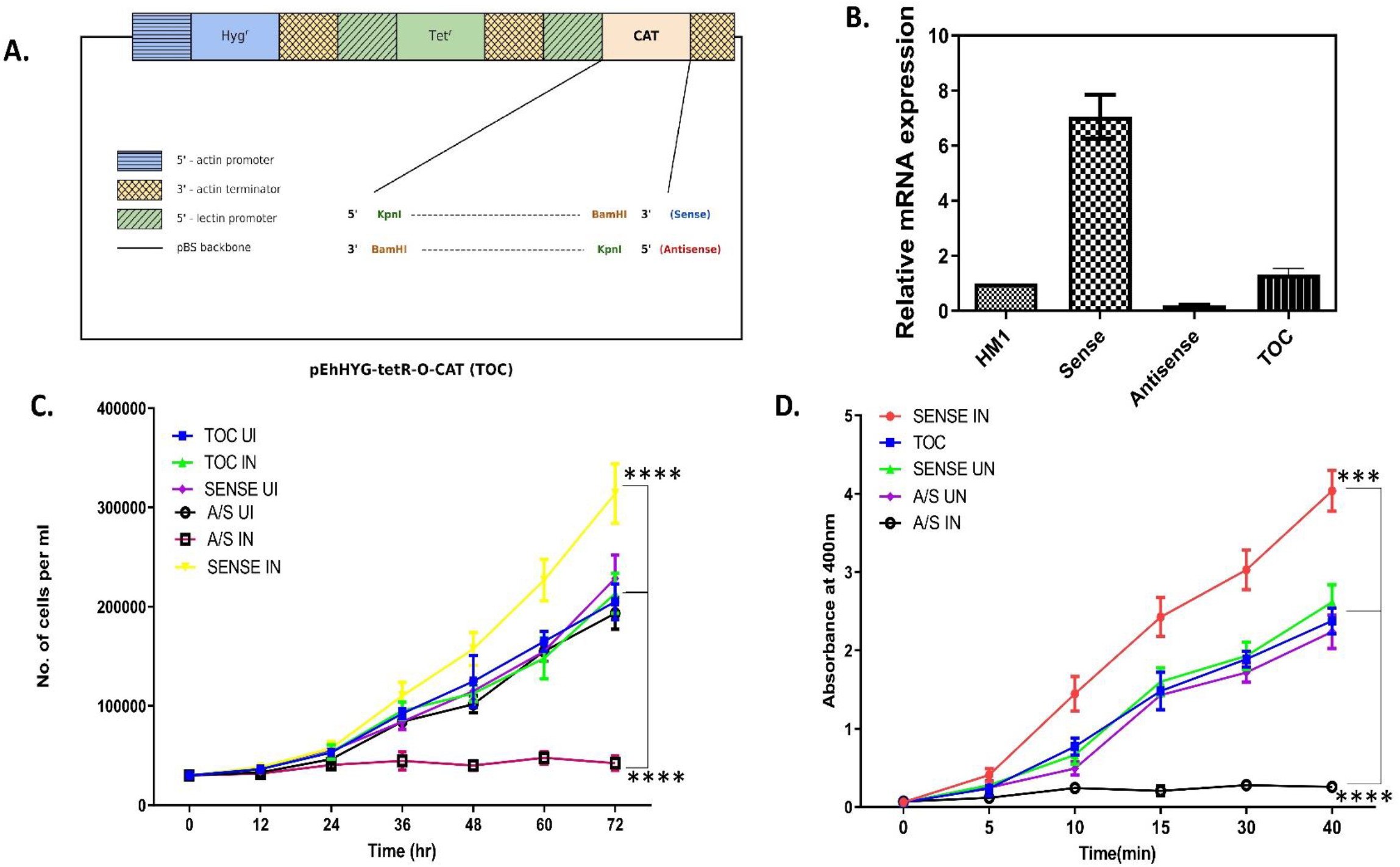
Effect of *Eh*PIG-K gene modulation on cell proliferation and phagocytosis in *Entamoeba histolytica*. **(A)** A schematic showing the tetracycline-inducible vector constructs (pEhHYG-tetR-O-CAT / TOC) which were designed for the expression of *Eh*PIG-K in the sense and antisense directions to upregulate and suppress the gene expression respectively. **(B)** The relative mRNA expression levels of *Eh*PIG-K in the wild-type (HM1), Sense *Eh*PIG-K (sense), Antisense *Eh*PIG-K (antisense), and vector control (TOC) cell lines as determined by qRT-PCR after induction. **(C)** The cell proliferation kinetics, expressed as the number of cells per mL, over a 72-hour period in induced (IN) and uninduced (UI) sense, antisense (A/S), and TOC control trophozoites. The cell line containing *Eh*PIG-K cloned in antisense direction shows 90% of growth inhibition in comparison to the control cell lines and cell line containing *EhPIG-K* cloned in sense direction. **(D)** The quantitative measurement of erythrophagocytosis (absorbance at 400 nm) over a 40-minute period in the induced and uninduced cell lines. The data are given as the mean ± SD from three independent experiments in duplicates (***p < 0.001, ****p < 0.0001). (Abbreviation: Sense UN- sense uninduced, Sense IN- sense induced, A/S UN- antisense uninduced, A/S IN – antisense induced, TOC- Tet-O-CAT vector alone, and HM1- wild type HM1: IMSS strain).

### 5. *Eh*PIG-K expression regulates the cell surface abundance of GPI-anchored Gal/GalNAc lectin

To assess the functional role of *Eh*PIG-K in GPI-anchor processing and in the presentation of the anchor in the membrane, we investigated the distribution and amount of Gal/GalNAc lectin, a major GPI-anchored surface protein in *Entamoeba histolytica* (Figure 5A). In trophozoites overexpressing *Eh*PIG-K (*Eh*PIG-K Sense), confocal immunofluorescence microscopy revealed strong and well-defined staining at the plasma membrane, while in those with *Eh*PIG-K suppressed (*Eh*PIG-K Antisense), the surface signal decreased markedly compared to the control (TOC) (Figure 5A). The mean fluorescence intensity (MFI) was used to show that the sense had a higher level of Gal/GalNAc lectin on the cell surface than the vector controls (TOC) (Figure B). On the other hand, the Antisense line showed very low surface expression, which indicates a severe reduction in the amount of the lectin at the plasma membrane (Figure 5B). An experiment involving parallel staining of the cell surface with ConA-FITC, bind various glycoproteins, glycolipids and α-D-glucose and a-D-mannose moieties in sugars, yielded similar results, showing greater surface labelling in the Sense line and a marked decrease in the surface signal after *Eh*PIG-K was suppressed (Supplementary Figure S2). Together, these findings show that the level of *Eh*PIG-K expression directly affects the amount and density of GPI-anchored surface proteins in *E. histolytica*.

**Figure 5:**
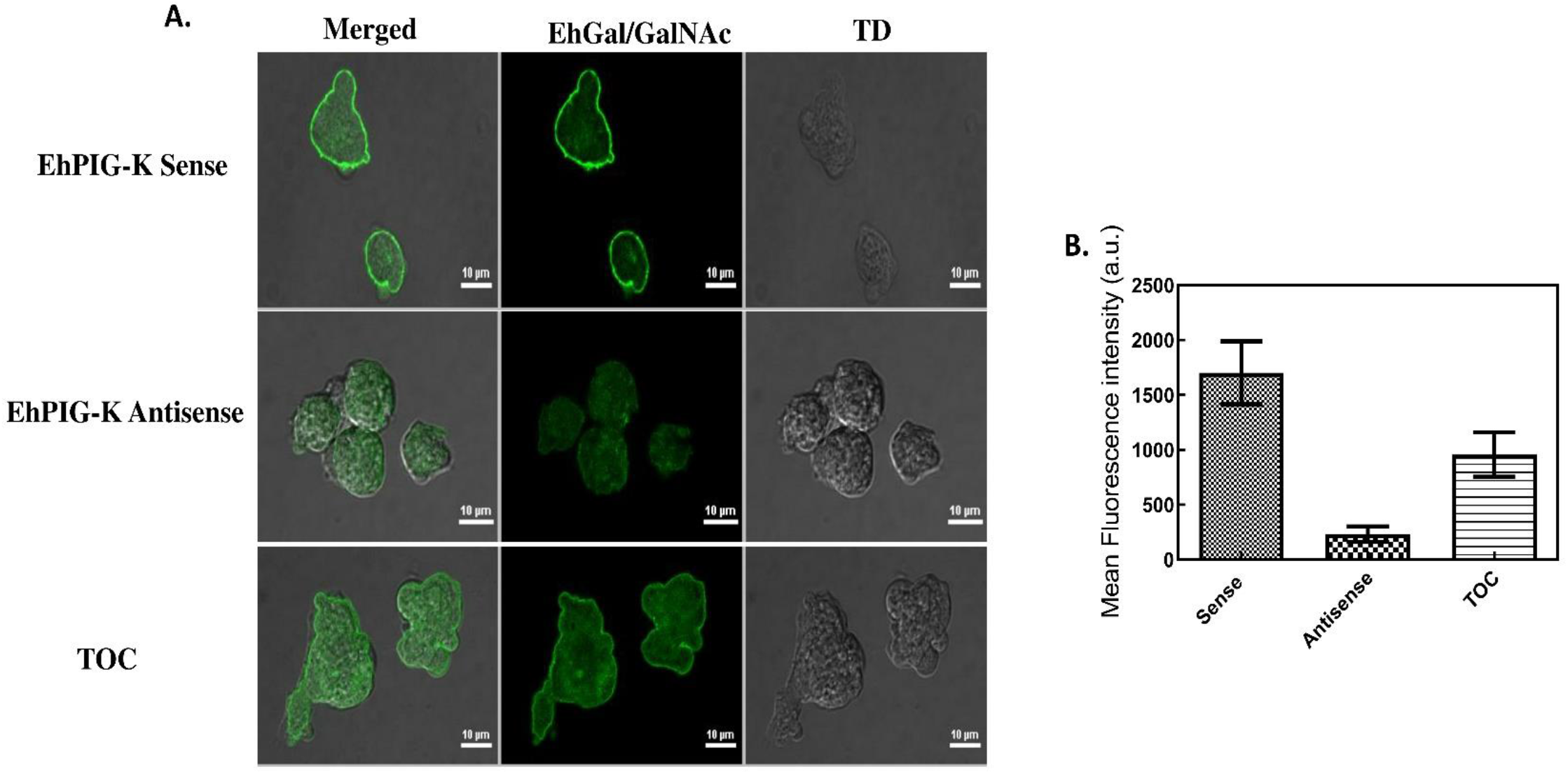
The levels of *Eh*PIG-K affect the expression of the Gal/GalNAc lectin on the cell surface in *Entamoeba histolytica*. **(A)** Confocal microscopy images obtained by immunofluorescence show the localisation of the Gal/GalNAc lectin (green fluorescence) on the cell surface in trophozoites that have overexpressed *Eh*PIG-K Sense, those in which *Eh*PIG- K has been suppressed *Eh*PIG-K Antisense, and those that are vector control (TOC). The panels show the merged image, the immunostaining for Gal/GalNAc, and the Transmission Detector (TD) brightfield channels. **(B)** The quantification of the Mean Fluorescence Intensity (MFI) in arbitrary units (a.u.), indicating the amount of cell surface lectin in the Sense, Antisense, and TOC control lines. The data are given as the mean ± SD of 50 cells from three independent experiments. (Abbreviation: a.u.- arbitrary units).

### 6. Subcellular distribution and Real-Time spatial localization of GFP tagged *Eh*PIG-K

To investigate the intracellular expression and spatial distribution of *Eh*PIG-K in *Entamoeba histolytica*, the GFP-fused constructs were examined using western blotting and confocal fluorescence microscopy. The expression of C-terminus *Eh*PIG-K-GFP construct was confirmed through western blot using anti-GFP antibody showed that the GFP-fused *Eh*PIG-K (33kDa *Eh*PIG-K +26 kDa GFP) was expressed in the total cell lysate (TCL lanes 1–4), compared with the vector control (GFP alone lane 6) (Figure 6A). Live-cell confocal imaging of *Eh*PIG-K-GFP cells under normal homeostasis revealed that the GFP-fused protein specifically localizes to distinct vesicular puncta (Figure 6B). Furthermore, these vesicular puncta were also observed during erythrophagocytosis, suggesting that the protein is indirectly involved in the dynamic cellular processes associated with phagocytosis. (Supplementary figure S3).

**Figure 6:**
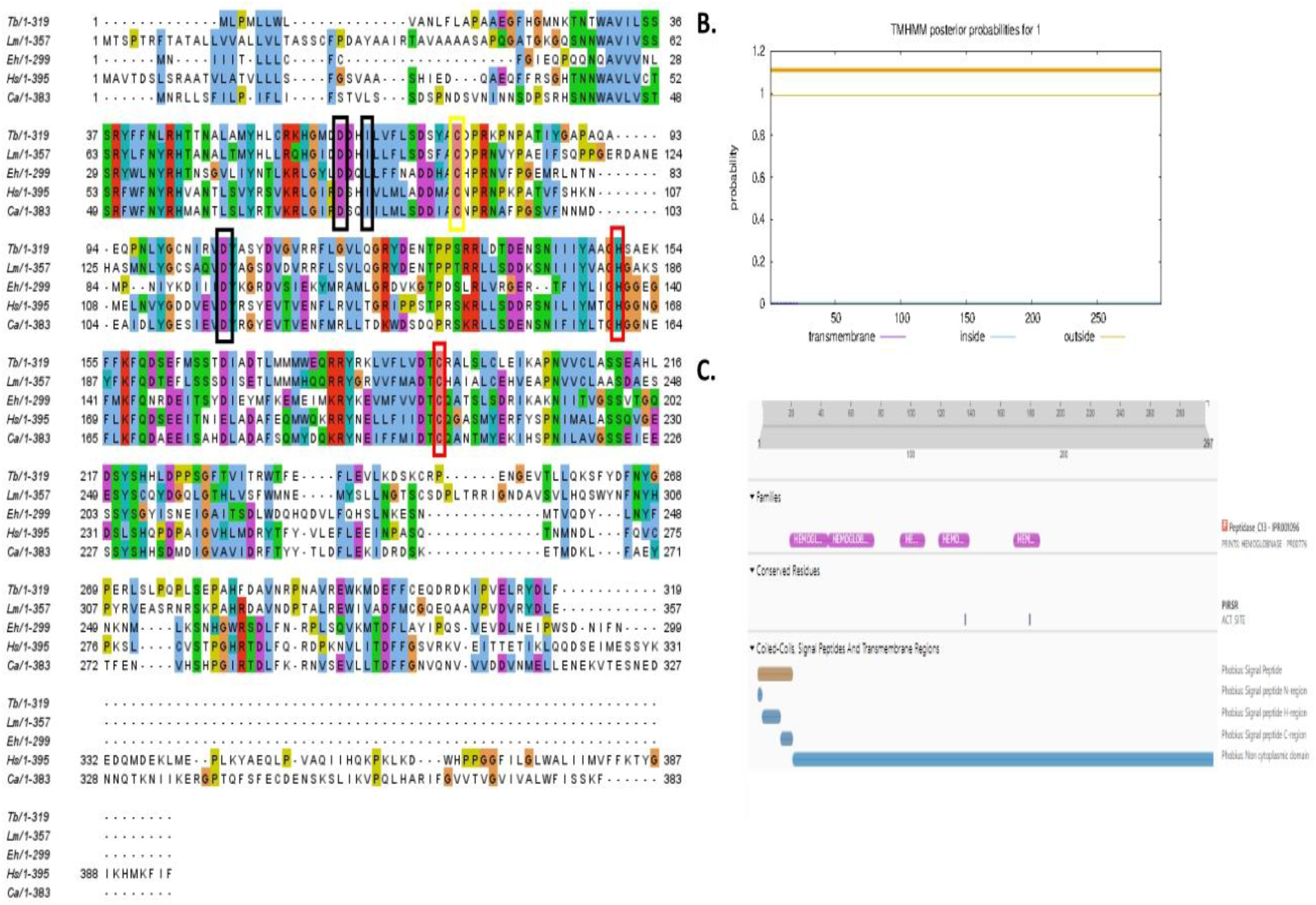
The subcellular distribution and real-time spatial localization of GFP-tagged *Eh*PIG-K. **(A)** Western blotting showing the expression pattern of GFP-tagged *Eh*PIG-K in the total cell lysate of *Eh*PIG-K-GFP transfected cell line induced with 30 μg/ml G418 (*Eh*PIG-k-GFP Lysate, lanes 1 to 4: 25 μg, 50 μg 75 μg, and 100 μg loaded respectively), lane 5 as HM1:IMSS control lysate (100 μg HM1 lysate) together with a GFP-only vector control lysate (lane 6: 100 μg only GFP lysate), and taking CaBP1 as a loading control. **(B)** Live-cell confocal microscopy showing the spatial localization of GFP-tagged *Eh*PIG-K in vesicular puncta alongside the corresponding transmitted light (TD) images.

### 7. In silico Sequence and Domain Architecture Analysis of *Eh*PIG-K

To assess the sequence characteristics and domain architecture of the *Eh*PIG-K, multiple sequence alignments and domain predictions were performed. When *Eh*PIG-K was subjected to multiple sequence alignment with its orthologs from *Trypanosoma brucei* (Tb), *Homo sapiens* (Hs), *Leishmania major* (Lm), and *Saccharomyces cerevisiae* (Ca), the catalytic residues and structural features were highly conserved. In particular, the catalytic dyad consisting of Cys178 and His136 (using *Eh*PIG-K numbering) is strictly conserved across all species (shown in red). Furthermore, an intersubunit disulfide bond formed by the conserved cysteine residues (highlighted in yellow) was detected, and Asp-55, Leu-58, and Asp-94 (using *Eh*PIG-K numbering) were identified as the metal-coordinating residues, indicating that the tertiary structure and functional integrity have been conserved (Figure 7A).

**Figure 7:**
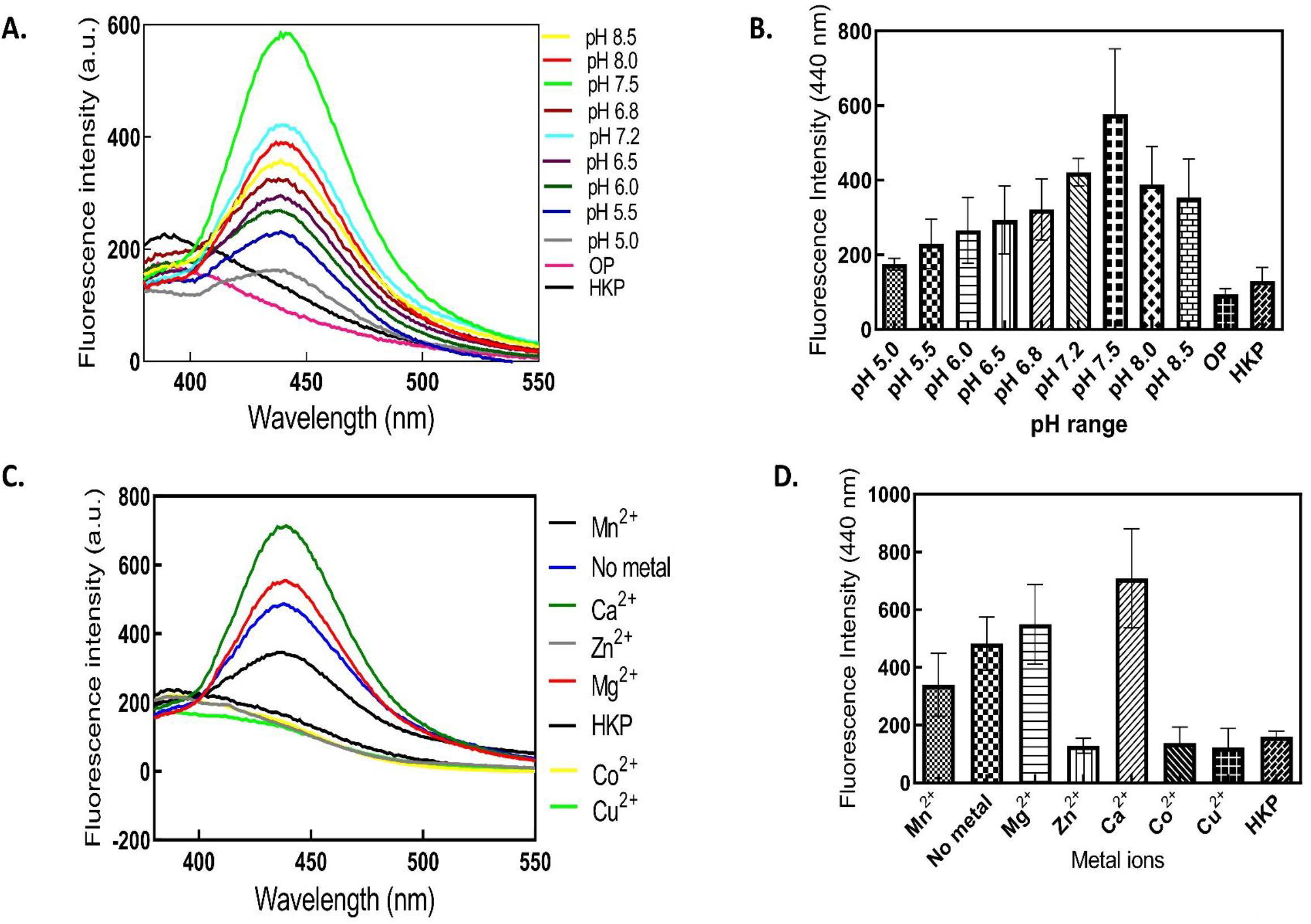
The sequence alignment, transmembrane topology, and domain structure of *Eh*PIG-K. A multiple sequence alignment using Clustal Omega is shown for *Entamoeba histolytica* PIG-K (*Eh*) together with its orthologs from *Trypanosoma brucei* (*Tb*), *Homo sapiens* (*Hs*), and *Saccharomyces cerevisiae* (*Sc*), highlighting the conserved catalytic dyad C178–H136 (indicated by red boxes), the cysteine residues (indicated by yellow boxes) which are involved in the formation of intersubunit disulfide bonds, and Asp-55, Leu-58, and Asp-94 were identified as the metal-coordinating residues **(A)**. Analysis of the transmembrane topology using TMHMM v2.0 indicates that the protein is entirely soluble/luminal (with an outside probability of 1.0) and has zero predicted transmembrane helices over the entire sequence length **(B).** Mapping of the domains by InterProScan reveals the conserved GPI-anchor transamidase / Peptidase C13 domain structure as well as the predicted N-terminal signal peptide and coiled-coil structural elements **(C)**.

Transmembrane region prediction was performed using the TMHMM v2.0 server. The posterior probability plot indicates that the entire protein sequence has a probability of 1.0 for the outside/extracellular (or luminal) orientation, with no predicted transmembrane helices along its length. This indicates that the submitted sequence corresponds to a soluble domain or a truncated/mature catalytic fragment lacking a hydrophobic membrane-spanning region (Figure 7B). Furthermore, the domain annotation by InterProScan indicated that *Eh*PIG-K has a conserved GPI-anchor transamidase domain and is a member of the Peptidase C13 (legumain-like) family (Figure 7C). The predicted signal peptide at the N-terminus, together with its transmembrane and catalytic features, supports its role in the GPI-anchor attachment pathway in *E. histolytica*. Unlike some higher eukaryotic GPI8 orthologs that have C-terminal hydrophobic transmembrane anchors, *Eh*PIG-K acts as a soluble or luminal-associated catalytic domain. This arrangement places its catalytic domain directly within the ER lumen, which corresponds to the orientation of the substrate processing sites on precursor proteins during transamidation.

### 8. In vitro peptidase assays demonstrate optimum pH and divalent cation activation of *Eh*PIG-K activity

In order to examine the catalytic properties of *Eh*PIG-K, the recombinant protein was purified using Ni^+2-^NTA affinity chromatography (Figure S1). The peptidase activity was measured using the fluorogenic peptide substrate ITTTNEN-AMC conjugated to AMC (designed from the Gal/GalNAc lectin). The assay was carried out in buffers of different pH and it was found that *Eh*PIG-K showed strong cleavage activity that was dependent on pH, the maximum relative fluorescence intensity being obtained at pH 7.5, whereas the activity was markedly decreased under acidic conditions (pH 5.0–6.0), and also under highly alkaline conditions. (Figure 8A and 8B). Additionally, when a range of divalent metal ions was tested, it was found that Ca^+2^ and Mg^+2^ significantly enhanced the catalytic activity, Mn^+2^ induced a minor increase in the activity, while ions such as Zn^+2^, Co^+2^, and Cu^+2^ inhibited peptidase activity as compared to the controls (Figure 8C and 8D). The heat-killed protein (HKP) and the substrate only controls (OP) were taken as negative controls, thus confirming the specificity of substrate cleavage by the active *Eh*PIG-K.

**Figure 8:**
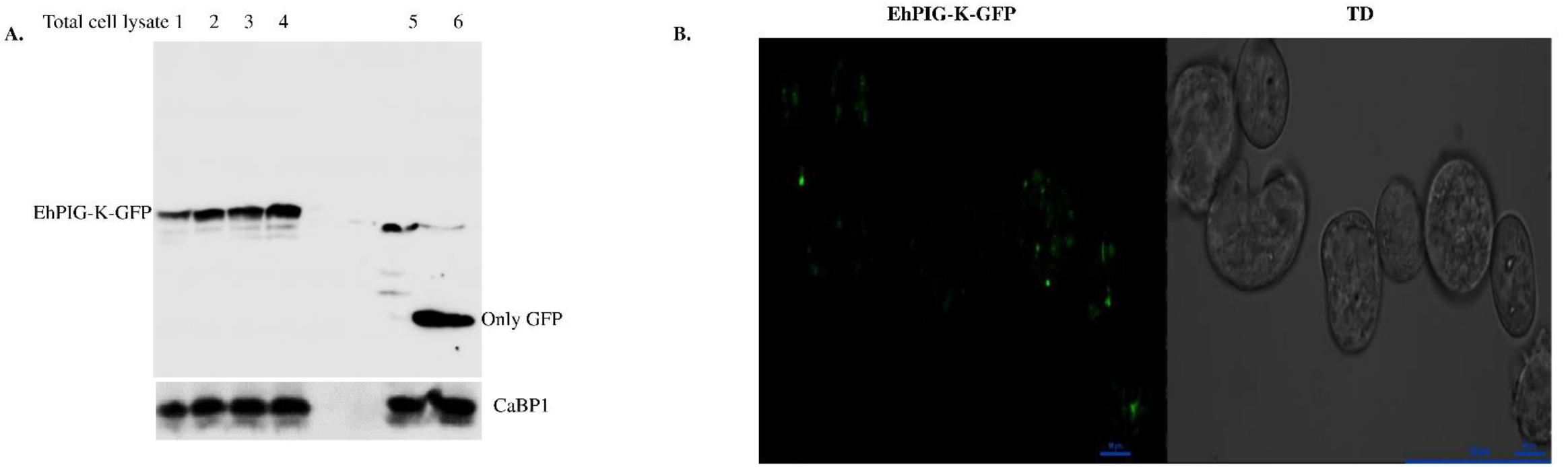
The enzymatic characterization and optimal conditions for *Eh*PIG-K peptidase activity. **(A)** The pH-dependent fluorescence intensity profiles and the related bar graph **(B)** indicate that *Eh*PIG-K showed peptidase activity best with the ITTTNEN-AMC substrate at pH 7.5. **(C)** The effect of different divalent cations on *Eh*PIG-K activity and the related bar graph **(D)** is shown, with Ca^+2^ and Mg^+2^ has highest increase in activity, Mn^+2^ induced a minor increase in the activity, while Zn^+2^, Co^+2^, and Cu^+2^ decrease it. Heat-killed protein (HKP) and substrate only (OP) were used as negative controls. The data show the mean emission at 440 nm. The purification profile is provided in Figure S1. The data are given as the mean ± SD from three independent experiments. (Abbreviation: a.u.- arbitrary units).

### 9. Preparation of the model of *Eh*PIG-K

All the models, including the Apo and the three ion-bound states (Ca^+2^, Mg^+2^ & Mn^+2^) of *Eh*PIG-K, were generated using the AlphaFold3 server. They were compared against the reference CryoEM structure of PIG-K in Human GPIT (PDB ID: 7WLD) and were found to fit well with an RMSD of around 1.0 [12]. The ion-bound states were also examined for the position of the ions. Structures were visualised and rendered using PyMOL (version 3.1.1, Schrodinger, LLC). We truncated the N-terminal disordered transmembrane region and prepared the soluble core of the protein harbouring the putative active site. For the molecular dynamic simulation, all the different states of the protein started from well-aligned initial structures (Figure 9A).

**Figure 9:**
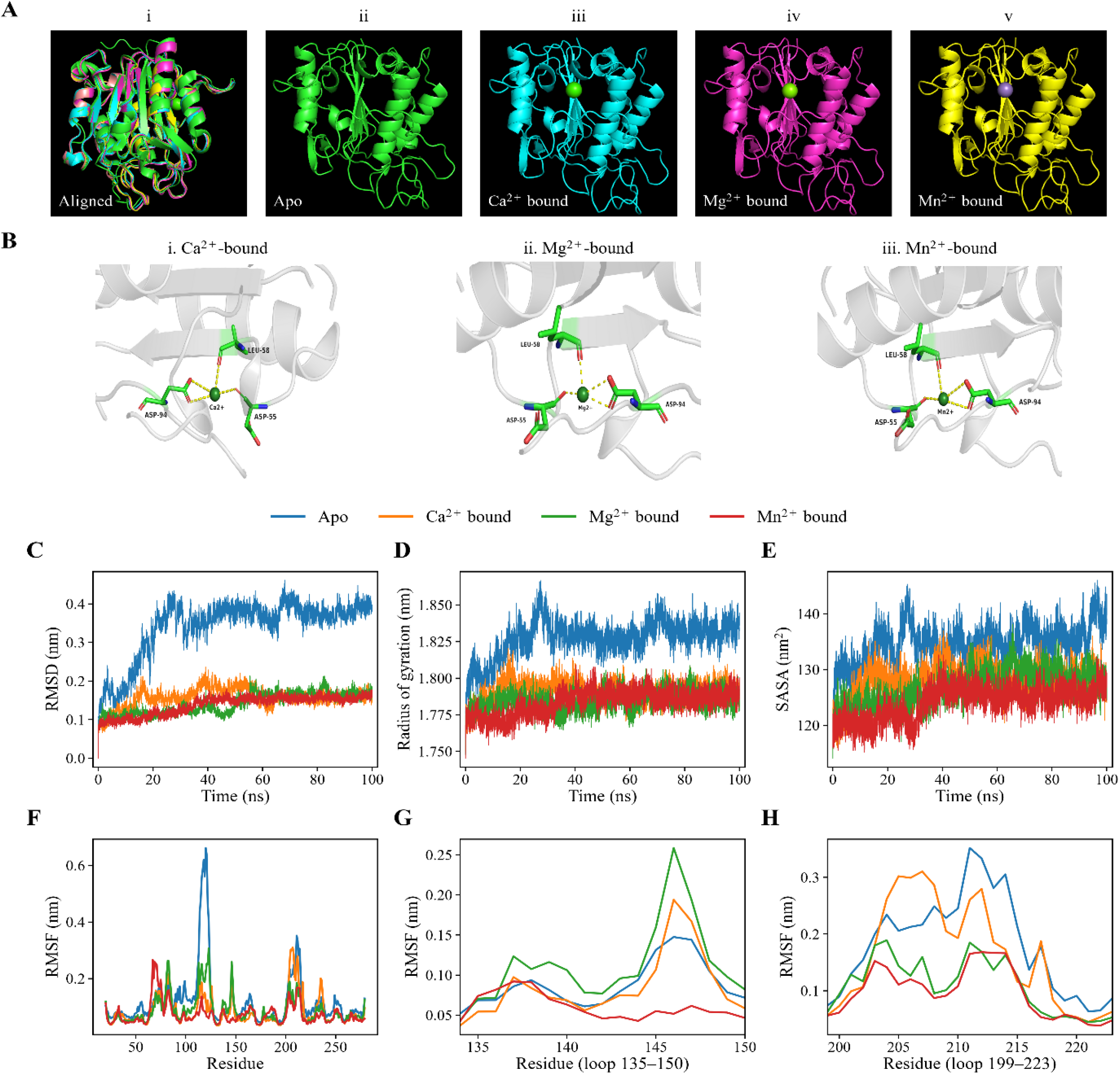
**(A.i-v)** *Eh*PIG-K initial models generated by AlphaFold3 server. Initial structures compared well amongst themselves, RMSD constantly being <1Å. (**B.i-iii)** Ion-coordinating residues in 3 ion-bound states of *Eh*PIG-K, namely, Ca^+2^, Mg^+2^, and Mn^+2^-bound states. (**C)** RMSD plot of 100 ns simulation of 4 states of *Eh*PIG-K. All the ion-bound states exhibits greater compaction of the structural core. (**D)** RG plot of 100 ns simulation of all the 4 states of *Eh*PIG-K. Apo *Eh*PIG-K differs from ion-bound states, but the difference is not substantial. (**E)** SASA plot of 100 ns simulation of all the 4 states of *Eh*PIG-K. Apo *Eh*PIG-K differs from the ion-bound states roughly by 7.4%. (**F)** RMSF plot of *Eh*PIG-K across the full-length of the protein. (**G)** RMSF plot of the region 132-150. (**H)** RMSF plot of the region 199-223. Figure (F), (G) and (H) suggest Mn^+2^-bound state restricts the local atomic fluctuations the most.

#### 9.1 Divalent ion binding leads to greater rigidity and compaction of the core domain

In all three metal-ion-bound forms, Ca^+2^, Mg^+2^, and Mn^+2^, three amino acid residues, namely Asp-55, Leu-58, and Asp-94, turned out to be the metal-coordinating residues. In *Eh*PIG-K, Leu-58 replaced the Ile participating in this coordinating triad as present in the other homologous proteins of other organisms, such as Asp-75, Ile-78, Asp-116 of *C. albicans* Gpi8, Asp-79, Ile-82, Asp-120 of human PIG-K, and Asp-63, Ile-66, Asp-106 of *T. brucei* PIG-K, respectively (Figure 9B.i-B.iii). The RMSD data calculated from the simulation run, plotted using Matplotlib, suggest that binding of the divalent metal ions such as Ca^+2^, Mg^+2^, and Mn^+2^ induces greater rigidity in the soluble core domain of the protein in comparison with the Apo form of the protein, which is more relaxed, with Mn^+2^ being the most effective amongst the divalent ions. The pattern was replicated in two out of three replicas (Figure 9C). Root-mean-square fluctuations (RMSF) indicate ion-dependent region-wise changes in local atomic fluctuations. The most prominent fluctuations were observed at or near the ion-coordinating region of the protein (loop 110-127 showed the greatest fluctuations in the Apo form). Binding of divalent cations pulls this region together and gives it a constant conformation (Figure 9F). There are two other regions of interest adjacent to the active site – (a) loop 135 – 150 that hosts the His136, which is part of the catalytic dyad of *Eh*PIG-K, and (b) the loop 199 - 223 that consists of many interacting residues that take part in positioning of the substrate peptide at the active site for optimum activity. Both regions exhibit ion-dependent differential behaviour, with Mn^+2^ being the most effective stabiliser of regional fluctuations (Figure 9G and H).

#### 9.2 SASA and RG analysis supports absence of any fold rearrangement of all the states

Solvent-Accessible Surface Area (SASA) analysis showing an upward deviation of the Apo *Eh*PIG-K structure from the other three metal-bound states, specifically around 30 ns, and the Apo form still wandering around 100 ns, suggests that the Apo form is more relaxed than the metal-bound states (Figure 9E). Radius of gyration (RG) analysis also supports the trend. The difference between the Apo and metal-bound forms in both plots is not substantial: RG differs by 0.04–0.05 nm, and SASA by roughly 7.4%, implying the absence of any major fold-rearrangement events (Figure 9F). From both analyses, we can credibly conclude that the Apo form is more expanded. At the same time, the metal ions pull the fold together more tightly, leading to greater compaction without any complete rearrangement of the fold.

#### 9.3 Energy calculations across all four states implies stable equilibrated system

The Lennard-Jones and potential energy plots showed stable fluctuations around a flat mean throughout the 100 ns simulation, indicating a well-equilibrated system with no sudden conformational fluctuations. That flat, overlapped Lennard-Jones plot across all four states suggests that the packing and vdW environments are essentially the same for all cases. (supplementary figure 4). The Mg^+2^ potential offset is not due to conformational changes but rather to other electrostatic or compositional terms.

#### 9.4 Active site-specific SASA rules out any noticeable difference in solvent accessibility of the active site surface

Active site SASA (supplementary figure 5A.i-A.ii) is comparable across the Apo, Ca^+2^, Mg^+2^ and Mn^+2^ simulations, indicating that the ion binding with the protein core does not substantially alter the overall solvent accessibility of the active site, so the activity differences based on ion-bound states are unlikely due to any significant changes in the active site exposure. While the inter residue distances plots between Cys178-Ser204 (supplementary figure 5B.iii) shows that Ca^+2^-bound state is maintaining a distance throughout the simulation, while the same is bringing the Thr217 somewhat closer to the Cys178 than the other forms (Figure 5B.iv). While the distance between Cys178 and Ser 218 and Cys 178 and Asp 219 are comparable amongst the ion-bound states and slightly lower than the Apo *Eh*PIG-K (supplementary figure 5B.v-B.vi). While there are not many differences in overall solvent accessibility of the active site surface, there are positional rearrangements amongst the residues that might have a say in the differential activity performance.

### 10. Cartesian Principal component analysis reveals localized variations in conformations of loops adjacent to the active site across the four states of *Eh*PIG-K

The RMSD indicated that the metal ions provide the protein soluble core greater rigidity. At the same time, RMSF indicated that Mn^+2^ is the constant stabilizer of local atomic fluctuations in both regions of interest (loop 135-150 & loop 199-223). SASA and RG analyses indicated comparable compaction across all four states. We wanted to investigate if there are any differential localized collective motions that may be influencing altered peptidase efficiency of the Apo and ion-bound states. To compare conformation sampling between the states, we performed a combined Cartesian principal component analysis across all 5 runs for each state. Out of the 780 eigenvectors, Principal component 1 appeared to explain 47.5% of the total variance in the global dominant collective mode; we focused our investigation on the top 6 principal modes, together explaining 68.5% of the total variance (supplementary figure S6A.i).

#### 10.1 Combined Cartesian principal component RMSF analysis revealed the contributions of local regions in the differential conformational ensemble of *Eh*PIG-K

To investigate the residues contributing most to the differing conformational ensembles along the dominant principal modes, we calculated Principal component analysis RMSF from the combined Cartesian PCA. RMSF analysis expanded the picture of contributions from regional common collective motions in the overall conformational differences. In the PC1 RMSF profile plot, the loop 110 – 128 is the dominant contributor along the PC1 mode. From the structural investigation of the 100 ns structure of each state (run 2), we can safely infer that in the Apo *Eh*PIG-K in the absence of ions, this loop is displaced (supplementary figure S6C.i), and that displacement dominated the PC1 mode. Study indicated towards two particular regions adjacent to the catalytic core comprising of Cys178 – (a) loop 135 - 150 harbouring the GHGG motif that is hosting the His136 of the catalytic dyad and G137, that is implicated to be partner in formation of the oxyanion hole alongside Cys206. Those residues are homologous to His164 and Gly165 of Human GPIT PIG-K subunit. And (b) loop 199 – 223, which consists of several functionally important residues like Ser203, Ser204, Ser206, Asp219, of them Ser 204 and Asp 219 are homologous to Human GPI-T PIG-K residues Ser232 and Asp247, which are part of substrate binding pocket S1 [37].

The loop 135 – 150 appeared to be particularly dominant in PC1, PC3, PC5, and PC6 modes, while the loop 199 – 223 appeared to be dominant in PC1 and PC2 modes (supplementary figure S6B.i-B.vi). For further investigation of the movements of both loops, we performed loop-specific Cartesian principal component analysis focusing on the loops 135-150 and 199-223.

#### 10.2 Loop-specific Cartesian principal component analysis reveals differential movement of loops adjacent to the active site among the states

We investigated the top 8 dominant common collective modes, which together explain 76.04% of the total variance, with the top 5 dominant modes explaining 62.75% of the total variance (supplementary figure S7A.i). In the loop-specific PCA, among the top 5 dominant modes, movement of the loop 135-150 dominates in PC2 and PC5; while movement of the loop 199-223 dominates in PC1, PC3, and PC4 (with a 1 Å threshold). We focused our investigation on the top 5 eigenmodes (supplementary figure S7B.i-B.v).

##### 10.2.1 Loop 135-150 conformational states depend on the state of the protein

Loop 135-150 hosts the “GHGG” motif, which contains two crucial residues: His136, that acts as a general acid, a crucial active site residue that is conserved in legumain as His148 and in Human GPIT PIG-K as His164 and oxyanion hole-forming partner Gly137 [38][13]. While the conformational ensembles of the loops along both modes are much more heterogeneous for the Apo *Eh*PIG-K across 5 simulation runs, the ion-specific movement patterns for the ion-bound states are replicated in at least 3 out of 5 individual simulation runs for each state. The Mn^+2^-bound state exhibits a very restricted conformational space compared to the Ca^+2^ and Mg^+2^- bound states in 4 of the 5 simulation runs (supplementary figure S8A.i-A.iv). In the GHGG motif region, although the exact position of His136 does not show any substantial movement in any of the states, the position of Gly137 shows a state-dependent differential pattern of movement.

Gibbs free energy landscape reflects the ion-dependent variations in the conformational ensemble in the common principal component space. Amongst the ion-bound states, the lowest-energy conformational basin for the Ca^+2^ and Mg^+2^-bound states sits at the negative PC2 regime compared to the Mn^+2^-bound states, while the space for the Mg^+2^-bound state explores a slightly more positive PC5 regime in contrast to the Ca^+2^-bound state, which is exploring both the regime. The Mn^+2^-bound state exhibits a closed conformational space, with the lowest-energy basin shifted into the negative PC5 regime compared to all other states of *Eh*PIG-K (Figure 10A.iv). While both the Ca^+2^ and Mg^+2^-bound states exhibit considerable movement of the loop (mean variance 6.582, 7.558 in PC1 mode, respectively) (supplementary figure S10 Table 1 and 2), they exhibit directional movement from the positive to the negative PC1 regime. While the Mg^+2^-bound state is a more active explorer of the space (supplementary figure S9A.ii-A.iii), the Ca^+2^-bound state samples from a broader conformational basin (Figure 10A.ii.). The aforementioned state-dependent differential movements of the loop 135-150 can be visually explained from the models along with both the modes (supplementary figure 10B and 10C). While the exact position of the His136 does not show much variation in movement among states, movement of Gly137 onwards may influence the active site geometry and electrostatic space of *Eh*PIG-K.

**Figure 10:**
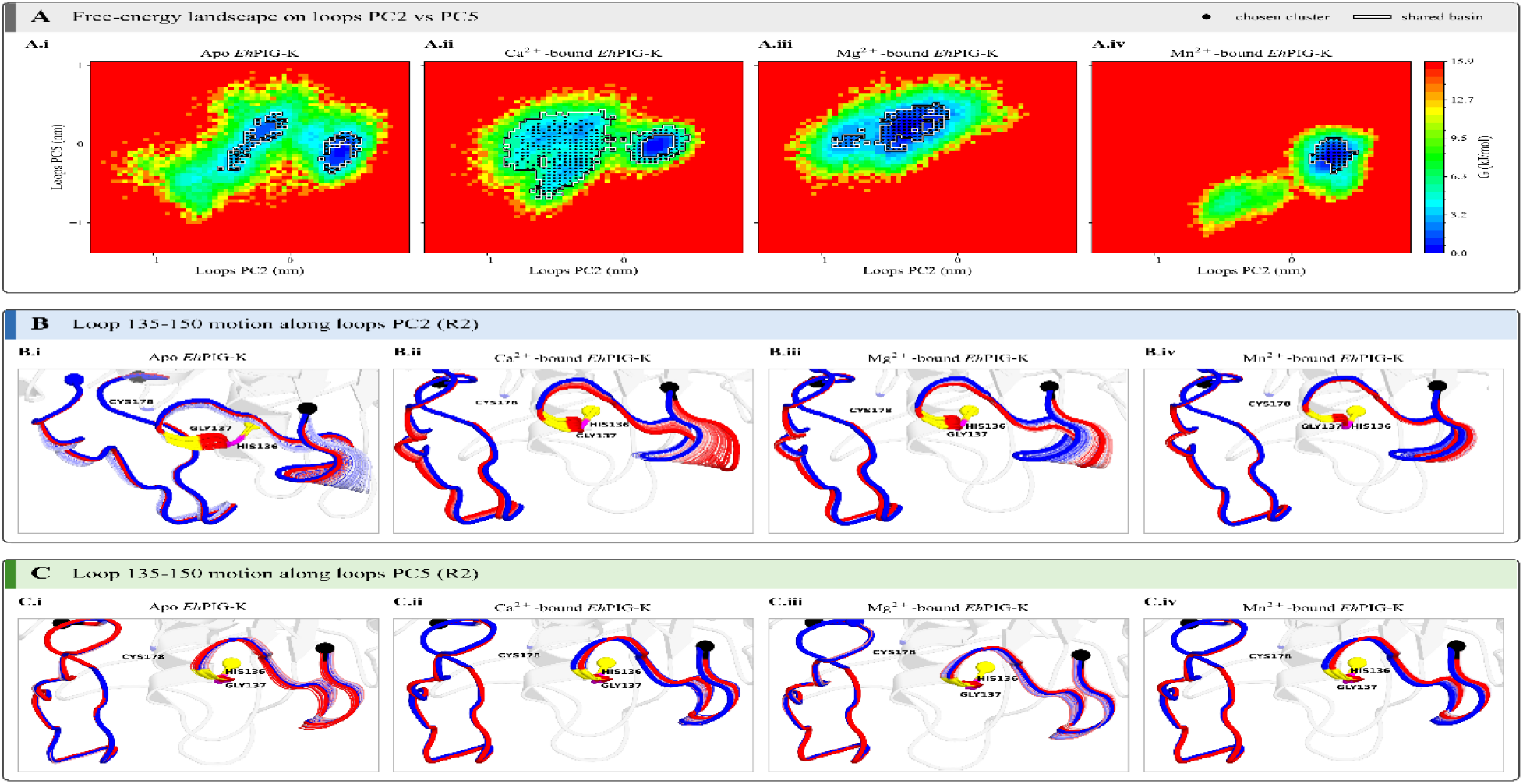
(**A)** Free energy landscape plots of PC2 vs PC5 common collective modes across all the 4 states. The loop conformation sampling is most broad and heterogenous in the Apo *Eh*PIG-K, same in the Ca^+2^ and Mg^+2^-bound state exhibits substantial movements, while the movement is lot more restricted and favoured conformational space is distinctively shifted towards positive PC2 and negative PC5 in Mn^+2^-bound state. (**B)** Movement of loops along the PC2 common collective mode. While Apo, Ca^+2^ and Mg^+2^-bound state exhibits substantial movement along the PC2 common collective mode, Mn^+2^-ion binding cripples the movement. (**C)** Movement of loops along the PC5 common collective mode.

##### 10.2.2 Ion-binding introduces altered conformational space and pattern of movement of loop 199-223

To investigate changes in the movement pattern of loop 199-223, which harbours substrate-interacting residues, we focused on the dominant common collective modes PC1, PC3, and PC4 as these modes are primarily dominated by the movements of loop 199-223. As proposed, Apo state of Human GPIT PIG-K stays dormant due to the auto-inhibitory positioning of loop-231(residue 231-237), with notable flip of Ser234, causing its Cα move by approximately 6Å to attain active state conformation(16). Our in silico study reveals there is an active state-dependent regulated movement of the loop 199-223 (corresponding loop of loop-231 of human GPIT PIG-K) that might be associated with differential activity of the states. Apo *Eh*PIG-K maintains a basal level of peptidase activity, which gets substantially higher on binding of Ca^+2^ and Mg^+2^ ions, with the Ca^+2^-bound state exhibiting the highest activity. While binding of Mn^+2^ significantly diminishes the activity. The regulated movement and positioning of the loop 199-223 might have the answer. Gibbs free energy reveals that while favoured conformational space of the Ca^+2^-bound *Eh*PIG-K sits around the same regime of the dominant common collective mode PC1 with the Mn^+2^-bound state, the conformational basin is much more restricted in case of the Mn^+2^-bound state. The projection of trajectory (supplementary figure S11) and time-resolved trajectory analysis shows that the Ca^+2^-bound state shows much more organized behaviour, sampling mostly from the 0 to negative PC1 regime while having occasional sampling from the positive PC1 regime (supplementary figure S13A.i-A.iv). In the Mg^+2^- bound *Eh*PIG-K, movement of the loop 199-223 is also heavily restricted as for the Mn^+2^- bound state (supplementary figure S11), while the favoured conformational basin is shifted to the positive PC1 regime (Figure 11A.iii). Both the Apo and Ca shows large movements along the PC1 direction (Figure 11C) (significantly higher mean variance and mean standard deviation, (supplementary figure S14 Table 1); the Apo *Eh*PIG-K shows a lot more heterogeneous movements, with frantic large excursions between PC1 regimes, in contrast to the Ca^+2^-bound state exhibits more organised behaviour (supplementary figure S13Ai).

**Figure 11:**
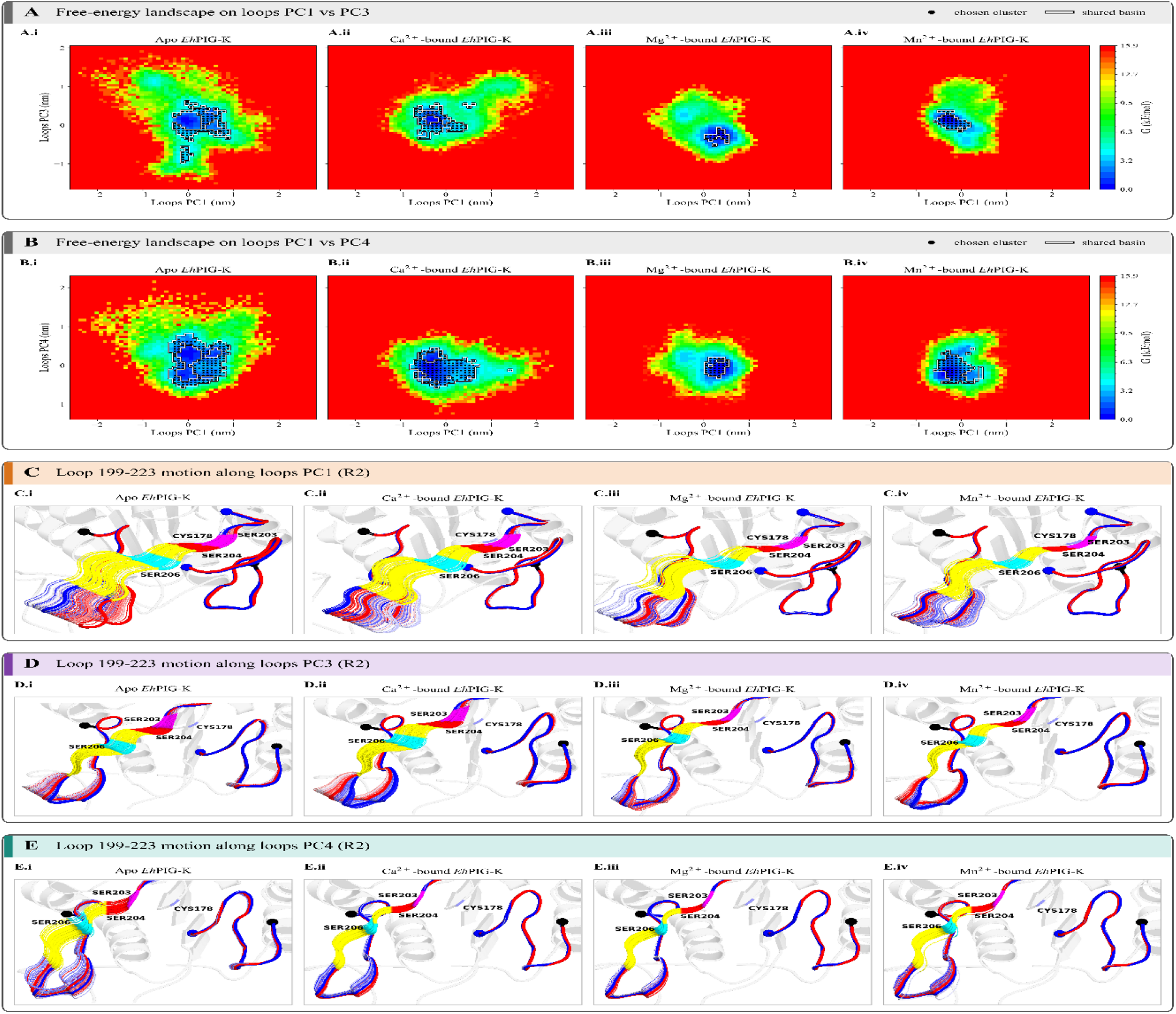
**(A)** Free energy landscape plots of PC1 vs PC3 common collective modes across all the 4 states. Apo *Eh*PIG-K exhibits heterogeneity in the movement, while Ca^+2^-bound state exhibits organised movement, while most favoured basin is shared in space with Mn^+2^-bound state, it relaxes the loop and allows sampling from less-favoured space. While Mg^+2^-bound state also shows restricted movement, the most favoured space is significantly shifted towards positive PC1 and negative PC3. (**B)** Free energy landscape plots of PC1 vs PC4 common collective modes across all the 4 states. All the three ion-bound states exhibit restrictions in the loop movement along PC4 common collective mode, Apo *Eh*PIG-K allows substantial movement. (**C)** Movement of loops along the PC1 common collective mode. (**D)** Movement of loop along the PC3 common collective mode. (**E)** Movement of loops along the PC4 common collective mode.

Along the PC3 common collective mode, though the loop 199-223 of the Ca^+2^-bound *Eh*PIG-K shows greater movement with respect to the other forms (mean variance of 13.529 Å) (supplementary figure S14 Table 2), followed by the Apo *Eh*PIG-K, it exhibits directional movements from the negative to positive PC3 regime (supplementary figure S13B.i-B.iv). In contrast, the Apo form shows extensive heterogeneity, with the Mg^+2^- and Mn^+2^-bound states being more restrictive (supplementary figure S11A.i-A.iv and supplementary figure S13B.i- B.iv). And in this mode either, although both in the case of Mg^+2^ and Mn^+2^-bound state the loop exhibits restricted movement (Figure 11D), the favoured conformational space is shifted to negative PC3 direction against the one of Mn^+2^-bound state (Figure 11Aiii).

Ion binding alters the pattern of the movement of loop 199-223 in both directions. The Ca^+2^- bound state allows significant motion of the loop; the Mg^+2^ and Mn^+2^-bound states restrict movement. A differential pattern is visible along the PC3 eigenmode, while the Ca^+2^-bound state allows the loop to move at par with the Apo state of the protein; the pattern of the movement is markedly different from the Apo state (supplementary figure 13B.ii). On the other hand, in the PC4 common collective mode, all of the ion-bound states show comparable restrictions in the loop movement, while more heterogeneous movement for the Apo state of *Eh*PIG-K (mean variance of 11.563Å) can be observed (supplementary figure S14 Table 3). Gibbs free energy landscape analysis exhibits a compact, equal-levelled, lowest-energy basin for all ion-bound states. In contrast, a more heterogeneous lowest-energy basin is observed for the Apo state of *Eh*PIG-K (Figure 11B) and can also be visualized from figure 11E.

### 11. Movement of the loops influences accessibility of the catalytic site *Eh*PIG-K

As we saw, the loop 199-223 exhibits extraordinary movement in both the Apo and Ca^+2^-bound form of *Eh*PIG-K, the loop 199-223 periodically closes in on the entrance of the catalytic site and drifts away from it, reopening the entrance (Figure 12B and 12C). That pattern looks a bit more unregulated in the case of the Apo, compared to the bit more periodic positioning in the case of the Ca^+2^-bound state. Both the Mg^+2^ and Mn^+2^-bound states exhibited restricted movement of the loop, but altered favoured conformational basins along the PC1 and PC3 common collective mode are observed to be effective in keeping the entrance to the catalytic site open through most of the time of the total simulation runs (Figure 12D). On the other hand, the Mn^+2^-bound state observed to keep Ser204 closed onto the entrance of the catalytic site, presumably barring the substrate peptide from getting optimum access to the catalytic site for catalytic efficiency; this might be the conformation of the inhibited state of *Eh*PIG-K (Figure 12E). The most favoured conformational basin of Ca^+2^-bound state also sits at the level of the Mn^+2^-bound state, while also allowing large excursions out of that basin, thus maintaining periodic access of the catalytic site while also maintaining a favoured geometrical space for catalysis.

**Figure 12:**
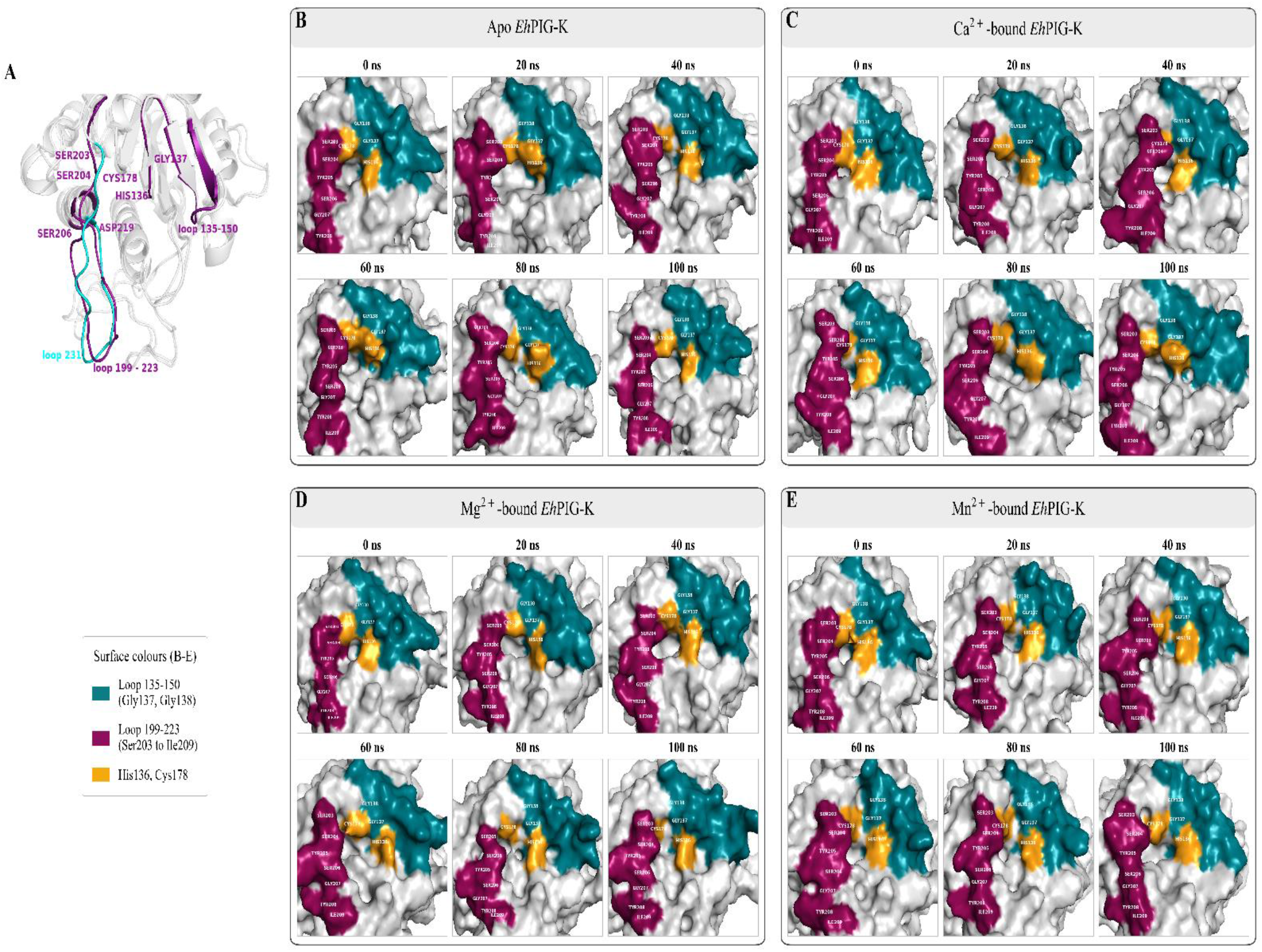
**(A)** Conserved residues between Human GPIT PIG-K and *Eh*PIG-K, important for its catalytic activity. (**B-D)** Positional alterations over the timeframe of the loop 135-150 and loop 199-223 across all 4 states. While the loop 199-223 in Apo *Eh*PIG-K closes in and drift away from the catalytic entrance in not so periodic manner, while in Ca^+2^-bound state the loop 199-223 seems to follow more periodic movement around the catalytic core. While both Mg^+2^ and Mn^+2^-bound state maintains restriction on the same, the shifted distinctively differed conformational space of Mg^+2^-bound state keeps the loop away from the entrance, thus keeping the entrance lot more exposed over the period, while the Mn^+2^-bound state moves keeps the loop residues close to the entrance of the catalytic residue.

### 12. State dependent variations in the movements of the loops adjacent to active site leads to differential distribution of the electrostatic potential across the active site surface

For the active site-specific analysis of the electrostatic surface, we selected four regions that collectively define the active site. Region 27-38 containing the residues – “NLSRYWLNYRHT”, region 132-150, that is covering the loop 135-150 prominently contributing to the common collective motion of the structural core, having the sequence – “LIGHGGEGFMKFQNRDEI”, region 174-179, that is harbouring the catalytic residue Cys-178, having the sequence – “VVDTCQ”, and the region 197-223, covering the loop 199-223, the other pronounced contributor to the common collective motion of the core, having the sequence – “SSVTGQSSYSGYISNEIGAITSDLWDQ”.

We observed that differences in the pattern of movements and favoured conformational spaces of the loops adjacent to the active site, along the dominant common collective modes, induce reorientation in the electrostatic surface distribution pattern of the active site. We can comfortably observe the state-dependent changes in the electrostatic potential distribution surrounding the deep negatively charged cleft, that is primarily the contribution of the residues 174-179, which hosts the catalytic residue Cys178 (Supplementary figure S15E). And that altered distributions of surface potential and the active site geometry end up positioning the peptide onto the active site (Figure 13A).

**Figure 13:**
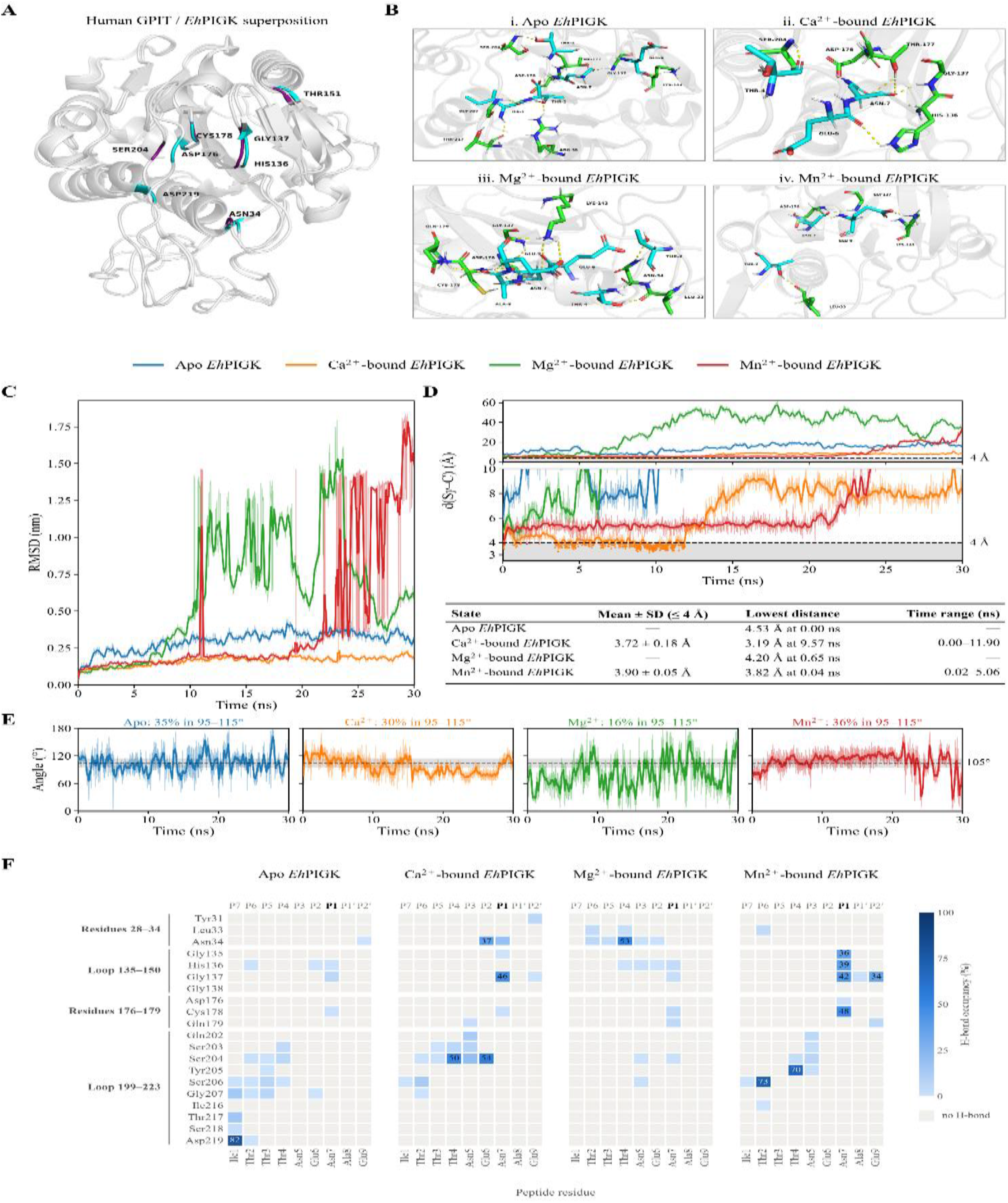
**(A)** Conserved residues between Human GPIT PIG-K and *Eh*PIG-K defining the active site of the proteins. (**B)** Interactions of the peptide sequence with the active site and adjacent residues of the receptor *Eh*PIG-K. (**C)** RMSD plot of the simulation of the receptor-peptide complex. (**D)** Distance between Cys178-Sγ of *Eh*PIG-K and carbonyl C of the scissile bond between Asn7 and Ala8. Ca^+2^-bound state exhibits greatest control, keeping the distance favourable for nucleophilic attack to happen. (**E)** Bürgi-Dunitz angle (α_bd_) plot of all four states of the simulation. All the states shows good maintenance of the angle. (**F)** Hydrogen bond occupancy heatmap of all four states of *Eh*PIG-K.

### 13. Differences in the interaction patterns of the peptide substrate with *Eh*PIG-K active site adjacent residues are observed

*Eh*PIG-K, being a cysteine protease of the C13 family (caspases and legumains are also members of the C13 family (citation), shares a good structural homology with another C13 family cysteine protease, Human GPIT PIG-K subunit. Essential residues, part of formation of the catalytic core including the catalytic dyad, trivalent oxyanion hole and substrate-binding pocket S1/S1’/S2’ were superimposed well with the Human PIG-K subunit of GPIT (PDB ID – 7WLD) (15). The active site residues Asn34, His136 and Cys178 are conserved against their homologous correspondences in Human GPIT PIG-K being Asn58, His164 and Cys206, and in legumains being Asn42, His148 and Cys189 (figure 13A) (18). Important residues of substrate-binding pockets for Human GPIT PIG-K – D204 and D247 of S1, G165 and Q207 of S1’ and T179A (16) were well matched with the corresponding residues in *Eh*PIG-K – D176, D219, G137, Q179 and T151 (Figure 13A). For positioning of the substrate, several combinations of restraints were checked to generate the best probable pose. Amongst them, the restraint combination having Asp176 OD1 or OD2 against Asn7 ND2 (H-bond donor), and Gly137 N of backbone N-H against O of carbonyl oxygen of scissile bond between Asn7 and Ala8 as part of probable oxyanion hole and Cys178 Sγ against the C of carbonyl carbon of scissile bond gave us the best performing pose.

100 ns MD-resolved structures were used for the docking, and state-dependent differential binding interactions were observed between the peptide and the active site residues (Figure 13 B). Altered orientation of the substrate against the active site can be observed from the different distribution of hydrogen bond occupancy heatmap (Figure 13F).

The Bürgi-Dunitz angle (α_bd_) of approach of the nucleophile to the C=O bond was established to be in the range 105+/-5⁰, later got refined to ⁓90⁰, for the successful nucleophilic attack. Here, all four states of the protein-substrate complex maintained the S_nuc_ … C_el_=O angle for some fractions of their complete trajectory, but their positional differences induced altered proximity of the Cys178-Sγ to the carbonyl C of the scissile bond. A favourable distance for a successful nucleophilic attack was efficiently maintained by the Ca^+2^-bound state (Figure 13D), a distance average of 3.72+/-0.18 Å, with the lowest distance being 3.19Å. Although we could not capture the favourable distance relationship for the Apo and Mg^+2^-bound state of *Eh*PIG-K, all these three states appeared to be ticking right boxes for a favourable positioning of the substrate for the peptidase activity. All the three states, namely Apo, Ca^+2^, and Mg^+2^-bound states transiently connect the residues D176, D219, G137, and Q179, which are catalytically important residues. Amongst them strong interaction of Gly137, coupled with transient H-bond interaction with Cys178, which are implicated as two oxyanion hole forming residues in the Human GPIT PIG-K and overall good connection with the Ser203 and Ser204, specifically evolutionarily conserved Ser204, analogous of which is Ser232, implicated to be part of S1 substrate binding pocket of Human GPIT PIG-K (16). For the case of the Ca^+2^-bound *Eh*PIG-K, may indicate a strongly favoured orientation of the peptide across substrate binding groove (Figure 13F). Both the Apo and Mg^+2^-bound state of the protein also maintain these crucial interactions, albeit transiently. While the Mn^+2^-bound state maintains some of these connections even more strongly, the strong distributional shifts in the connections imply a different orientation of the peptide across the active site. It may be the result of an altered conformation of the active site due to strict closed positioning of the loop 199-223, which might have its influence in lowered activity performances.

## Discussion and conclusion

The glycosylphosphatidylinositol (GPI) anchoring pathway is one of the most important post- translational modification systems in eukaryotes, as it enables attachment of a wide variety of proteins to the extracellular leaflet of the plasma membrane [39][40]. In protozoan parasites, GPI-anchored proteins (GPI-APs) make up a major group of cell-surfacemacromolecules that function as the main mediators of pathogenesis by causing adherence to tissues, enabling invasion of host cells, allowing the parasite to evade the immune response, and facilitating the acquisition of nutrients [39]. A multi-protein subunit GPI transamidase complex carries out the cleavage of the C-terminal signal peptide from precursor proteins and, at the same time, transfers the preassembled GPI core lipid precursor to the newly formed ω-site [9]. The catalytic subunit PIG-K/GPI8, which is a conserved cysteine endopeptidase [41], plays a key role in this process. In the human intestinal parasite *Entamoeba histolytica*, the major virulence factors such as the Gal/GalNAc lectin complex and PPGs—depend entirely on GPI anchoring for their display on the membrane and for their function [4][5]. Although its clinical significance is well recognized, the biochemical nature, intracellular behaviour, and exact role of the catalytic transamidase subunit, *Eh*PIG-K, in controlling parasite physiology have, until now, remained largely unknown.

The multiple sequence alignment in silico domain analysis showed that *Eh*PIG-K has the characteristic structural features of the Peptidase C13 (legumain-like) family of cysteine proteases. When the sequences of the eukaryotic orthologs it was found that the catalytic dyad made up of Cys178 and His136 is conserved and this pair consisting of a nucleophilic cysteine and a basic histidine forms a catalytic dyad mechanism which is necessary for the cleavage of the peptide bond at the ω-site and for the subsequent formation of a thioester intermediate before the phosphoethanolamine group of the GPI preassembled anchor carries out a nucleophilic attack [12]. Moreover, the sequence alignment identified a conserved cysteine (highlighted in yellow) that takes part in an important intersubunit disulfide bond with PIG-T, helping to maintain the tertiary structural integrity and the stability of the complex that is needed for assembly with non-catalytic transamidase subunits such as PIG-S, PIG-T, PIG-U, and GPAA1 [42]. Topological predictions made with TMHMM v2.0 indicated a probability score of 1.0 for being extracellular or luminal, and no transmembrane helices were predicted. The InterProScan domain mapping confirmed the presence of an N-terminal signal peptide which directs the newly formed polypeptide into the lumen of the endoplasmic reticulum (ER).

In order to assess the enzymatic activity of *Eh*PIG-K, a fluorogenic peptidase assay was performed using a synthetic peptide substrate (ITTTNEN-AMC), where AMC act as a reporter and has previously been used to study the peptidase activity of GPI8 [43] [44]. In *Trypanosoma*, it was shown that recombinant GPI8 purified from *E.coli* was active *in vitro* [43]. The purified *Eh*PIG-K showed strong and dose-dependent peptidase activity, thus providing direct experimental evidence that *Eh*PIG-K is a catalytically active enzyme that is able to recognize and cleave the GPI substrate sequences. The enzymatic characteristics revealed a clear sensitivity to pH, with maximum cleavage occurring at pH 7.5. The activity fell sharply under acidic conditions (pH 5.0–6.0) and decreased under highly alkaline conditions. This optimum pH value, which is physiological, corresponds exactly to the neutral to slightly basic pH environment that exists in the ER lumen. Metal ions enhance the peptidase activity; *Candida* GPI8 activity is metal ion dependent caspases, and activity is enhanced by Mn^+2^ [44], where as human GPI8 is Ca^+2^ dependent [12]. Analysis of our results showed that divalent ions, namely Ca²⁺ and Mg²⁺, greatly increased the endopeptidase activity of *Eh*PIG-K, whereas Mn²⁺ had a lesser increase.

*Eh*PIG-K is located within the intracellular ER network and does not appear on the cell surface. Interestingly, the ER structure observed was highly dependent on the fixative method used. When the cells were fixed with 3% paraformaldehyde (PFA), a fine, punctate, vesicular pattern was obtained because protein cross-linking may occur more slowly or may be less extensive, and during this fixation process, the ER network may undergo structural rearrangement, collapse, or fragmentation before it is fully immobilized. Consequently, the ER may appear fragmented or vesicular rather than as a continuous network. While fixation with 4% PFA resulted in an extensive reticular network spread throughout the perinuclear area and beneath the plasma membrane. A quantitative analysis revealed a strong spatial correlation with the ER Tracker and with the well-known luminal ER resident marker Calreticulin, also known to be localised in the plasma membrane during phagocytosis and bind to the human c1q [33]. It is also known to be involved in phagocytosis of apoptosis and cancer cells [45] [46]. Most importantly, immunofluorescence experiments carried out on intact (non-permeabilised) and detergent-permeabilised trophozoites proved that *Eh*PIG-K does not localise to the external surface of the plasma membrane but co-localised internally with EhCRT and Gal/GalNAc lectin in the ER regions. The low correlation observed with nuclear DAPI staining further confirmed that *Eh*PIG-K is located in the ER reticular network and in the continuous nuclear envelope membrane, without accumulating in the nucleoplasm. It was reported in *Drosophila* that GPI8 is located in the ER, while other subunits of GPIT complex are located in the ER as well as the nuclear envelope [47]. In *Trypanosoma*, this GPI8 subunit is localized in the tubular ER network [43].

In *Entamoeba histolytica,* protein trafficking and intracellular compartmentation are considerably different from the usual eukaryotic patterns. The intracellular endomembrane system—made up of the ER, Golgi-like structures and dynamic vesicular networks—forms an interconnected continuum rather than consisting of separate compartments [48] [49]. In order to check whether *Eh*PIG-K undergoes dynamic vesicular trafficking into degradation or endocytic pathways during active membrane remodelling, we examined its distribution during erythrophagocytosis. Even though the endomembrane network is fluid, *Eh*PIG-K is excluded from late endosomes that are positive for Rab7a and from vacuoles that are positive for LysoTracker. In a physiological context, this lack of presence in acidic compartments means that *Eh*PIG-K does not undergo unwanted proteolytic degradation by lysosomal hydrolases ruling out the endo-lysosomal pathway as a destination for *Eh*PIG-K and strongly supports its function as a specialized protein that resides in the ER where GPI anchor attachment takes place to the nascent proteins prior to their anterograde transport through the Golgi-like compartments towards the cell surface.

There are a few studies of functional characterization of *E. histolytica* GPI biosynthetic pathway genes using antisense RNA [6] [7]. When the growth of the parasite was monitored over a period of 72 hours, a marked difference in their proliferation was observed. When *Eh*PIG-K was suppressed, it caused reduced cell growth and significantly inhibited trophozoite proliferation in a time-dependent manner. Suppression of *Eh*PIG-K disrupts the function of the GPI transamidase complex, leading to the accumulation of unprocessed and unanchored protein precursors within the endoplasmic reticulum (ER) lumen. This accumulation induces severe ER proteostatic stress and arrests cell cycle progression. As a result, the absence of newly synthesized GPI-anchored proteins (GPI-APs) at the outer membrane causes surface starvation, signalling failure, and subsequent population decline. [50][51][52]. At the same time, normal parasite survival and division depend on a steady level of surface expression of GPI-anchored receptors, nutrient transporters, and signal transducers [53]. Just as severe growth defects and cell death have been observed after disruption of the transamidase gene in other parasitic protozoa, in *Trypanosoma brucei*, a conditional knockout of *GPI8* results in rapid cell cycle arrest and the accumulation of uncleaved variant surface glycoprotein (VSG) precursors, ultimately leading to cell death [14]. Similarly, *Leishmania mexicana* mutants that lack GPI8 show a loss of surface leishmanolysin (gp63) and have impaired amastigote survival [54]. These comparative results highlight the conserved essentiality of the PIG-K subunit among divergent eukaryotic human pathogens.

The main clinical feature of infection with *E. histolytica* is invasion and destruction of tissue, a process which is caused by adherence to host cells and erythrophagocytosis [55][56]. The Gal/GalNAc lectin is a major GPI-anchored surface heterodimer and is responsible for binding to the mucin of the host colon, to epithelial cells and to erythrocytes [57]. The direct relationship between *Eh*PIG-K expression and surface lectin density demonstrates that transamidase levels act as a rate-limiting step in the maturation and delivery of GPI-anchored virulence factors in *E. histolytica*. Since the uptake of erythrocytes depends on recognition mediated by the Gal/GalNAc lectin and on contact-dependent wrapping of the membrane [58], the absence of surface lectins in the suppressed cells means that the trophozoites are unable to adhere to the target red blood cells [6]. Mechanistically, unanchored lectin precursors that do not undergo transamidation are held back by the internal ER quality-control system and are directed for proteasomal degradation, thus preventing their transport to the plasma membrane.

Our in-silico investigation for the reason of the differential kinetic efficiency from structural point of view indicates state-dependent altered pattern of movements of two loops adjacemt to the catalytic site of *Eh*PIG-K might have a say in its catalysis. RMSD, RG, and SASA analyses revealed that the presence of metal ions led to greater compaction of the protein core. At the same time, the not-so-substantially different and stable RG and SASA profiles for the Apo form of *Eh*PIG-K suggest a perceived relaxation in the sampling probability of the Apo conformational ensemble. Still, the amplitude of the difference between RG and SASA indicates that events such as unfolding and refolding of the protein core are absent. On the other hand, the active-site-specific SASA suggests no substantial changes in the solvent accessibility of the active-site pocket, implying that the kinetic difference may not result from differential exposure of the active-site pocket.

We assumed the difference in the conformational state was due to localised differential behaviour, which led us to analyse combined Cartesian PC RMSF, that expanded the image. State-dependent differential movement of the loop 135 – 150 harbouring the His-136 residue in its GHGG motif, part of the catalytic dyad of the protein, and Gly-137, partner in formation of oxyanion hole, and the loop 199 – 223 hosting several of the substrate interacting residues of the substrate binding pockets S1/S1’/S2 that help in the positioning of the substrate in the active site pocket with respect to the Cys-178.

Loop 135-150 exhibits substantial movement along both the PC2 and PC5 common collective mode in Apo, Ca^+2^ and Mg^+2^-ion bound states. While the conformation of the loop in Ca^+2^- bound state samples from a slightly overlapping space with Mg^+2^-bound state, favoured conformational space is significantly restricted and shifted to positive PC1 and negative PC5, distinctively different from the other two ion-bound states. And in the Apo *Eh*PIG-K this loop exhibits lot more heterogenous movement along both the PCs. While the exact position of His136 doesn’t take part in any substantial movement, the state-dependent differences in the pattern of the movement of the region of Gly137, which itself is part of formation of oxyanion hole, and onwards results in differential differed orientation of active site geometry and electrostatic potential distribution around the catalytic entrance.

As in Human GPI-T PIG-K a flipping mechanism of the residues Ser232, Ser234 and His 235 send the loop-231 in more active conformation from the closed autoinhibited state, our study on *Eh*PIG-K reveals a state-dependent movement of the loop 199-223 (corresponding loop of loop-231of Human GPI-T PIG-K) may be the reason of the differential peptidase activity among the states. In Apo *Eh*PIG-K loop 199-223 moves in a heterogenous and frantic manner, and controls the exposure of the entrance to the nucleophilic core, but in an unregulated manner. Ca^+2^ ion binding to the protein provides the movement of the loop a structured manner, the loop controls the entrance to the nucleophilic Cys178 in much more periodic manner, closes in towards the active site groove at some points, and release out subsequently. This way it may control the entry of the peptide, subsequent positioning of it and also influence the interatomic connections required for the activity better. On the other hand, in the other two ion-bound states (Mg^+2^ and Mn^+2^-bound states), the movement is lot more restricted in a closed space, but the relative positioning of the loop differs amongst them. Loop 199-223 in Mg^+2^-bound state stays positioned away from the entrance to the catalytic residue Cys178 for most of the timespan, thus keeping the entrance exposed for the peptide. But the lack of movement may result in failure to maintain the optimum positioning of the peptide with respect to the catalytically important residues, thus leading to the reduced peptidase efficiency. On the other side, the positioning of the loop 199-223 in Mn^+2^-bound state is significantly shifted towards the entrance to the catalytic residue Cys178. Though the most favoured conformational basin Ca^+2^- bound state shares the space with Mn^+2^-bound *Eh*PIG-K, the Ca^+2^-bound states relaxes the loop, let it sample from less favoured spaces, thus maintaining the geometry of the active site and entrance to the catalytic nucleophilic agent in regulated manner, while Mn^+2^-bound state fixes the loop in closed-in position towards the entrance. That may act like a guard, inhibiting the entry of the peptide to the catalytic core and also influence a suboptimal placement of it for the peptidase activity to happen, thus leading to a very low peptidase efficiency.

All these state-dependent differences in the collective movements of the loops adjacent to the active site pocket in the dominant eigenmodes also lead to altered electrostatic distributions within the active site pocket, that leads to the altered positioning and orientation of the substrate peptide with respect to the nucleophilic agent Cys178-Sγ and oxyanion hole-forming components – amide N of Gly137 and Cys178. That results in the differential peptidase capability amongst the states of *Eh*PIG-K.

In summary, our findings indicate that *Eh*PIG-K is a cysteine peptidase localized to the endoplasmic reticulum. It is essential for GPI anchor processing, cell-surface presentation of the major virulence factors, proliferation, and erythrophagocytosis in *Entamoeba histolytica*. Upon genetic suppression of *Eh*PIG-K, the essential surface adhesins are lost, caused a lethal arrest of cell proliferation. It shows that the GPI transamidation machinery represents a vulnerable bottleneck for parasite viability. Therefore, *Eh*PIG-K provides a highly novel and promising target for drugs against parasites. Small-molecule inhibitors that target the cysteine dyad in the active site of *Eh*PIG-K could selectively inhibit the expression of the parasite’s surface virulence factors, resulting in the death of the parasite and offering a promising approach for the treatment of amoebiasis.

## Acknowledgments

The authors acknowledge the Central Instrument Facility, School of Life Sciences, JNU, for support with confocal microscopy and fluorescence spectroscopy. From the same school, we sincerely thank Prof. Sneha Sudha Komath for her insightful feedback and manuscript editing, and Dr. Shalini Mishra for her assistance with initial culture handling.

## Author Contributions Statement (CRediT)

Simra: Executed all aspects of the study, including conceptualization, methodology, experimentation, analysis, and preparation of the original draft as well as review and editing.

Arnab Das: Conducted in silico studies and contributed to manuscript preparation. Zenab Kamali: Assisted with methodolody.

Prof. S. Gourinath (corresponding author): Supervised all aspects of the study, including conceptualization, methodology, project oversight, and manuscript review and editing.

## Ethics Statement

All procedures involving animals were conducted with the national guidelines established by the Committee for Control and Supervision of Experiments on Animals (CPCSEA), which operates under the Ministry of Environment, Forest and Climate Change, Government of India.

