## Supplementary Data for "Characterization of *Eh*PIG-K, a catalytic subunit of the GPI transamidase complex in *Entamoeba histolytica*"

**Table S1** lists all primers used in the study.

| | |
|---|---|
| SENSE_PIGKFOR | 5' CGGGTACCATGAACATCATCATTACCTTGCTT 3' |
| SENSE_PIGKREV | 5' CGGGATCCTTAATTAAAAATGTTATCTGACC 3' |
| ANTISENSE_PIGKFOR | 5' CGGGATCC ATGAACATCATCATTACCTTGCTT 3' |
| ANTISENSE_PIGKREV | CGGGTACC TTAATTAAAAATGTTATCTGACC 3' |
| FOR_PIGK_TOC_FLAG | 5' CGGGTACCATGAACATCATCATTACCTTGCTTTTGTGTTTTTGTTTTGG 3' |
| REV_PIGK_TOC_FLAG | 5' CGGGATCC TTACTTGTCGTCATCGTCTTTGTAGTCCTTGTCGTCATCGTCTTTGTAGTC ATTAAAAATGTTATCTGACC 3' |
| FP_GFPPIGK_KPN1_CGFP | GGGGTACCCCATGAACATCATCATTACCTTGCTT |
| RP_GFPPIGK_ECOR5_CGFP | GGATATCATTAAAAATGTTATCTGACCAAGGTATT |
| FOR_PIGK | CCATGGGCATGGTTGTCGTTAATTTATCAAGATA |
| RP_PIGK | 5' CCGCTCGAGATTAAAAATGTTATCTGACCAAGGTATT 3' |

**Table S2** list of primers used for Real-time quantitative reverse transcriptase-PCR

| | |
|---|---|
| EhPIG-K FP | TTAATGCTGACGACCACGCA |
| EhPIG-K RP | TCTCTTCCAAGCATTGCTCTCA |
| 18S rRNA forward primer | 5' CGCCCGTCGCTCCTACCG 3' |
| 18S rRNA reverse primer | 5' CTTTTAATGATCCTTCCGCAGG 3' |

### Supplementary Figure S1

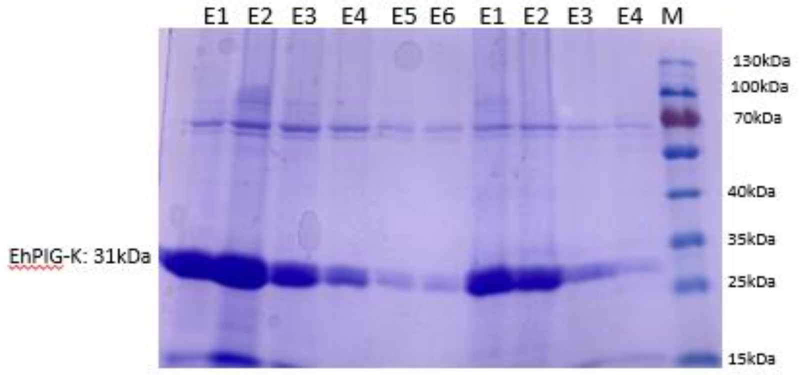


**Figure S1**. Purification of Recombinant *Eh*PIG-K: Recombinant *Eh*PIG-K was expressed in *E. coli* and purified using $Ni^{2+}$-NTA affinity chromatography. SDS-PAGE analysis of the elution fractions (E1–E6) showed a homogenous band at the expected molecular weight of ~31 kDa. Lane M represents the molecular weight marker.

### Supplementary Figure S2

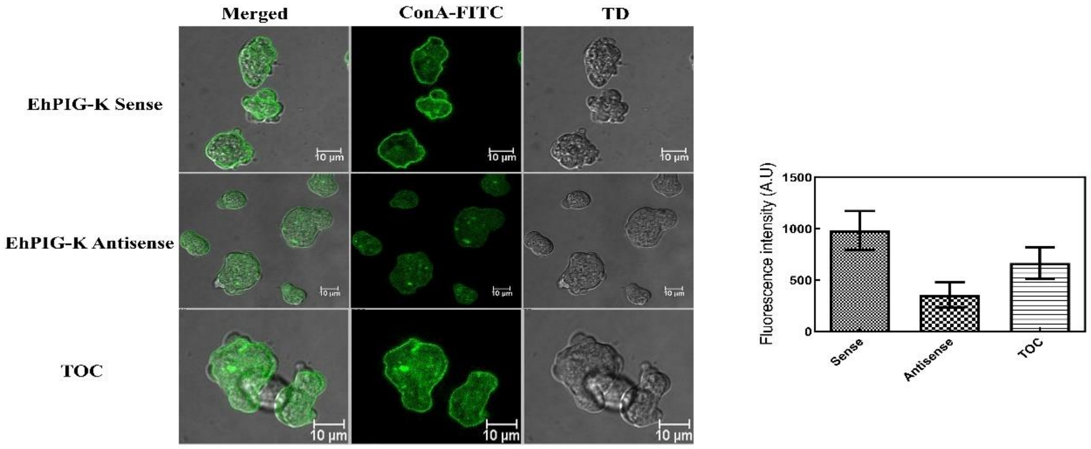


**Figure S2**: The levels of *Eh*PIG-K affect the binding of the ConA-FITC on the cell surface in *Entamoeba histolytica*. **(A)** Confocal microscopy images obtained by immunofluorescence show the binding of the ConA-FITC (green fluorescence) on the cell surface in trophozoites that have overexpressed *Eh*PIG-K (Sense), those in which *Eh*PIG-K has been suppressed (Antisense), and those that are vector control (TOC). The panels show the merged image, the ConA-FITC staining, and the Transmission Detector (TD) brightfield channels. **(B)** The quantification of the Mean Fluorescence Intensity (MFI) in arbitrary units (a.u.), indicating the amount of cell surface lectin in the Sense, Antisense, and TOC control lines. The data are given as the mean ± SD from 50 cells.

**Supplemenraty Figure S3**

**Figure S3:** Live cell imaging movie of GFP-*Eh*PIG-K during erythrophagocytosis, suggesting that the protein is localised in the vesicular puncta and indirectly involved in the dynamic cellular processes associated with phagocytosis.

**Supplementary Figure S4:**

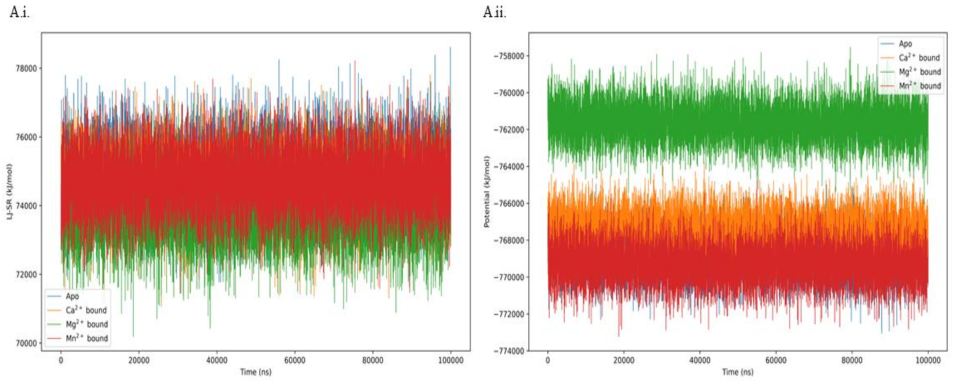


**Figure S4: (A.i)** Leonard-Jones (short-range) energy plot for all four states. (**A.ii)** Potential energy plot for all four states

**Supplementary Figure S5:**

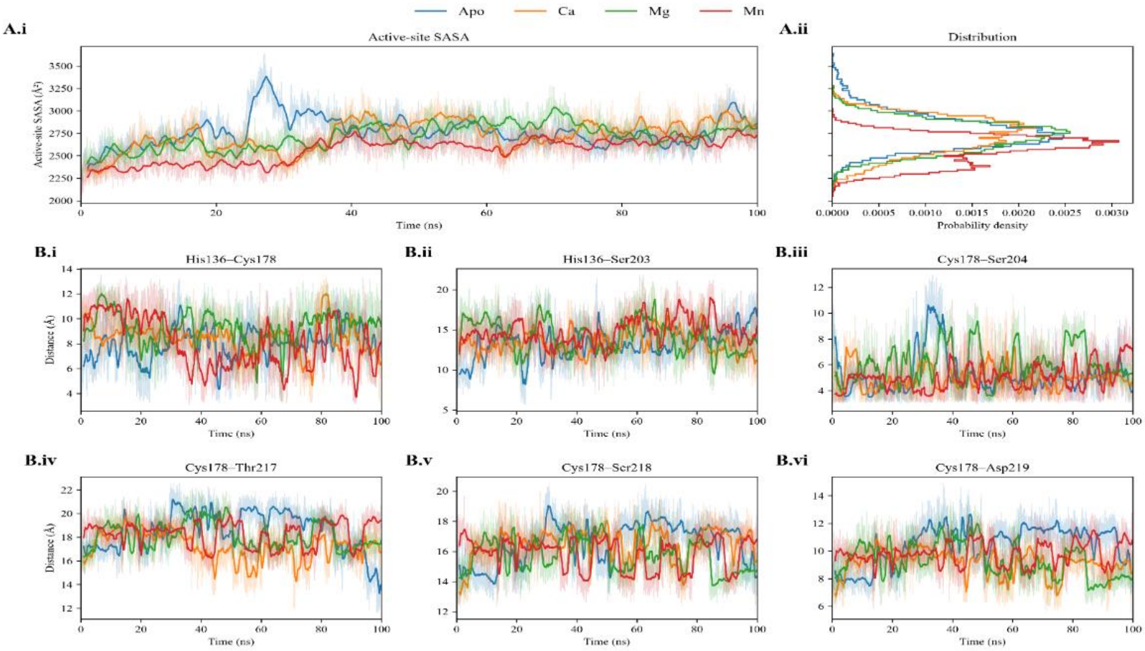


**Figure S5: (A.i)** Active site SASA plot across the all four states. (**A.ii)** Probability distribution across the all four states. (**B.i-B.vi)** Inter-residue distance between active site adjacent residues and the partners of catalytic dyad – Cys-178 and His-136.

**Supplementary figure S6:**

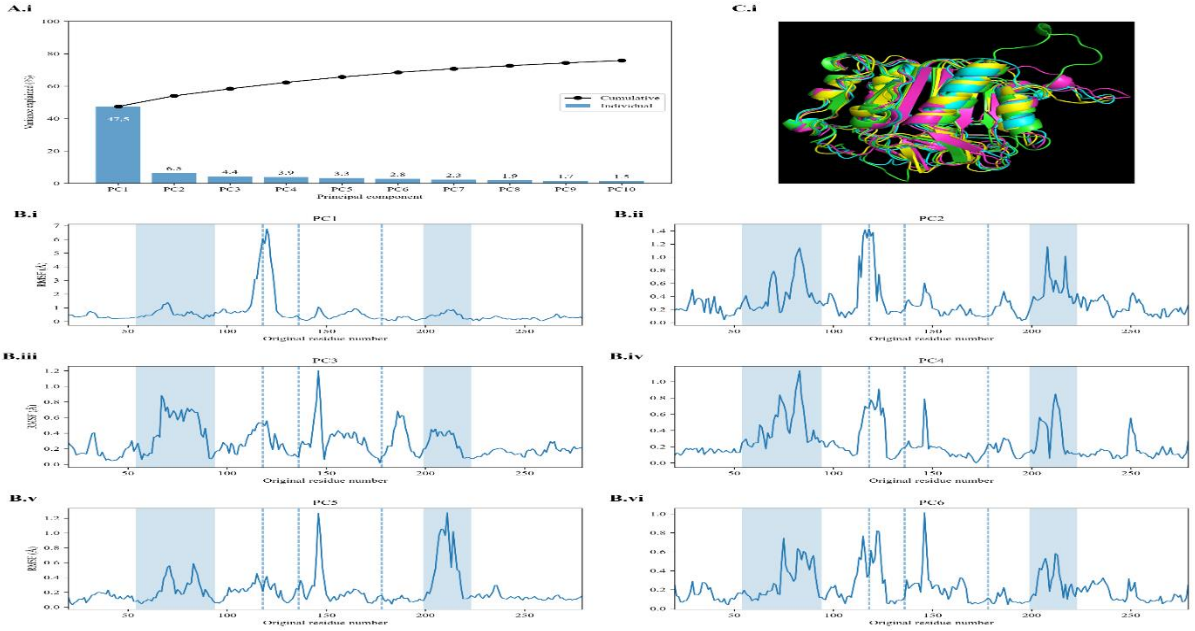


**Figure S6: (A.i)** Variance explained by the top 10 eigenvectors of the combined whole protein backbone Cartesian principal component analysis. (**B.i-B.vi)** combined PC RMSF along top 6 eigenvectors. (**C.i)** Displaced loop 110-128.

**Supplementary figure S7:**

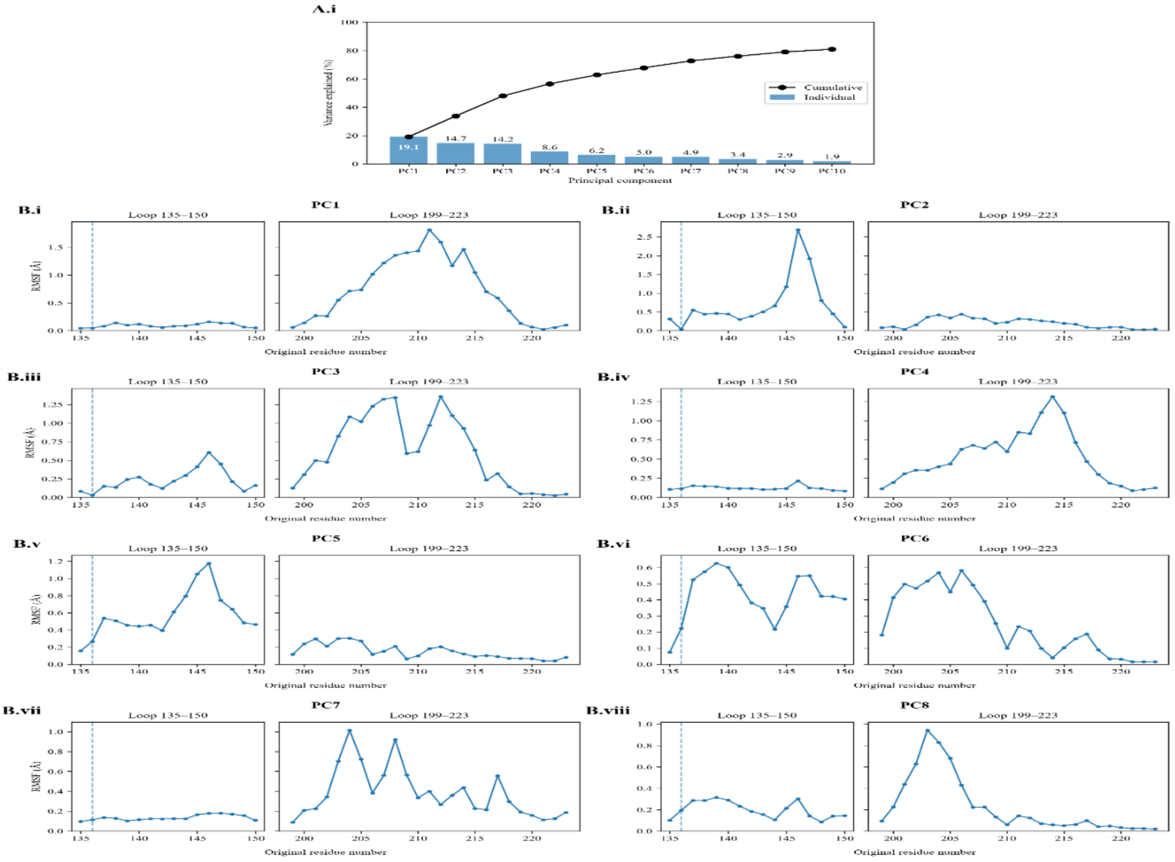

**Figure S7: (A.i)** Variance explained by the top 10 eigenvectors of the loop-specific combined Cartesian principal component analysis. **(B.i-B.viii)** loop-specific combined PC RMSF along top 8 eigenvectors.

### Supplementary figure S8:

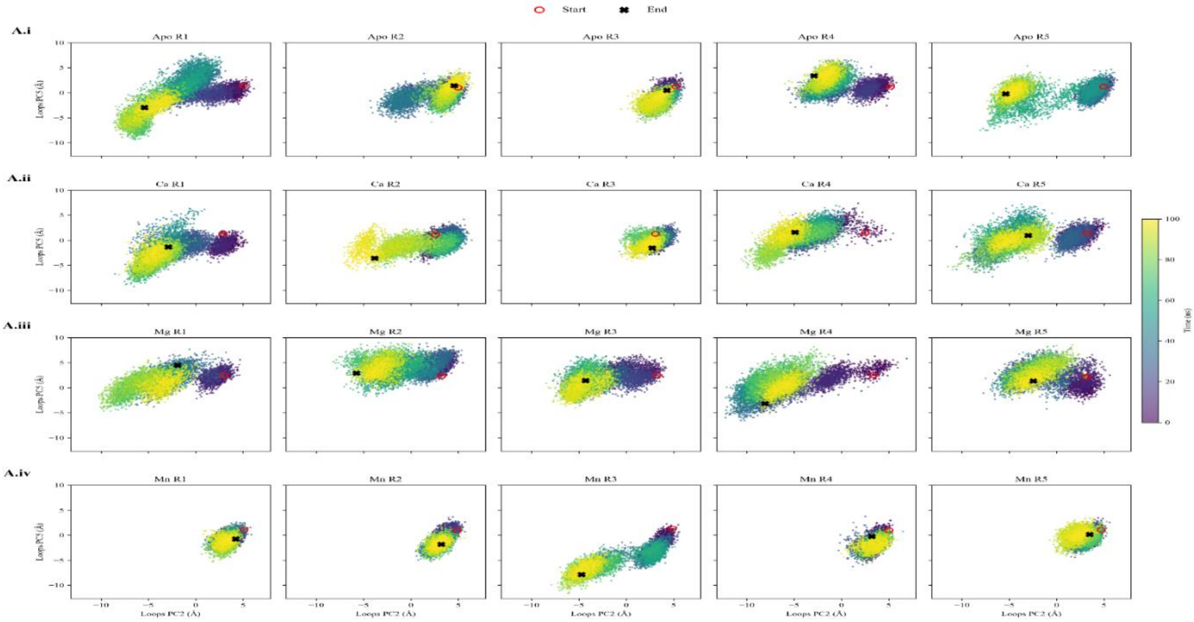


**Figure S8: (A.i-A.iv)** Projections of the trajectories of 5 runs of each state of loop 135-150 backbone (C-α atoms) onto the PC2 and PC5 eigenvector components of loop-specific combined Cartesian PCA.

### Supplementary figure S9:

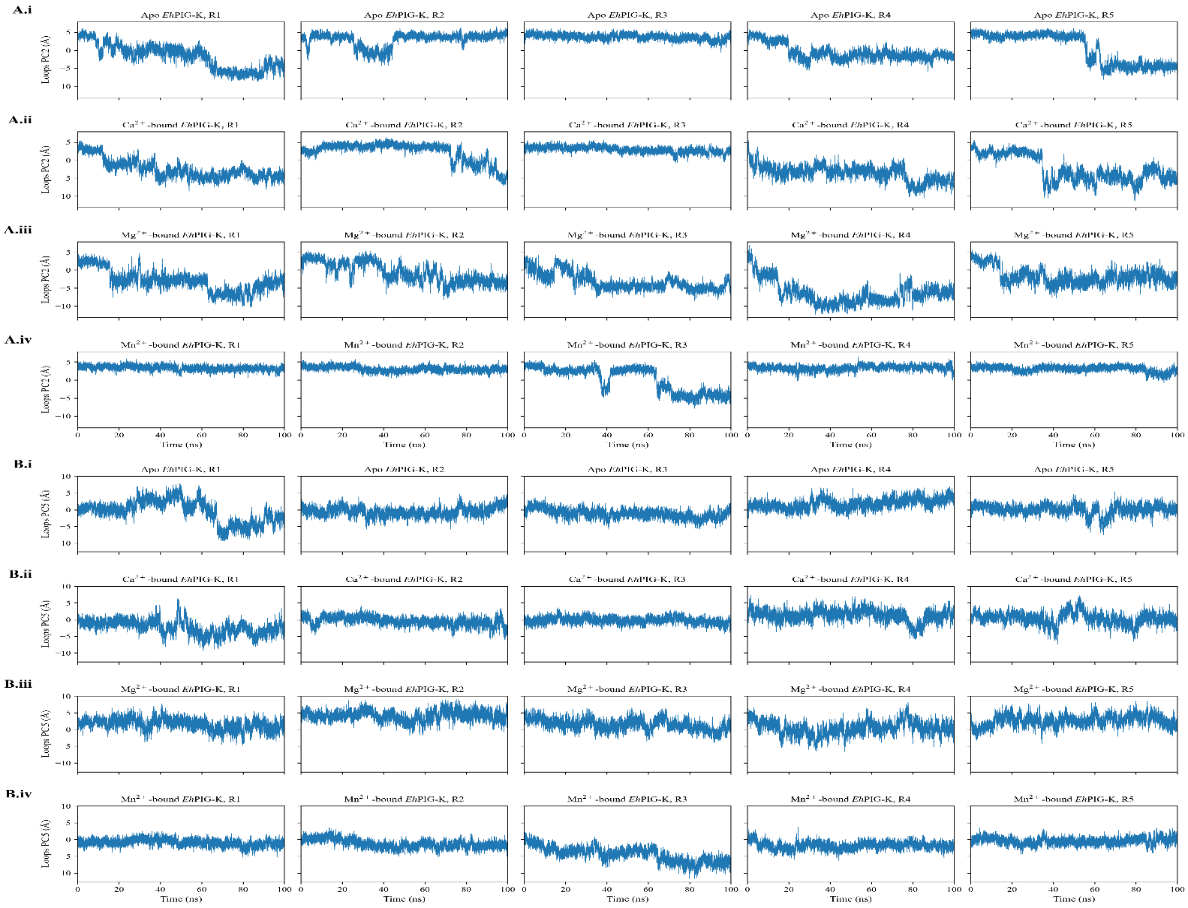

**Figure S9: (A.i-A.iv)**Time-resolved projections of loop 135-150 backbone trajectories onto PC2 eigenvector components. (**B.i-B.iv)** Time-resolved projections of loop 135-150 backbone trajectories onto PC5 eigenvector components.

### Supplementary figure S10:

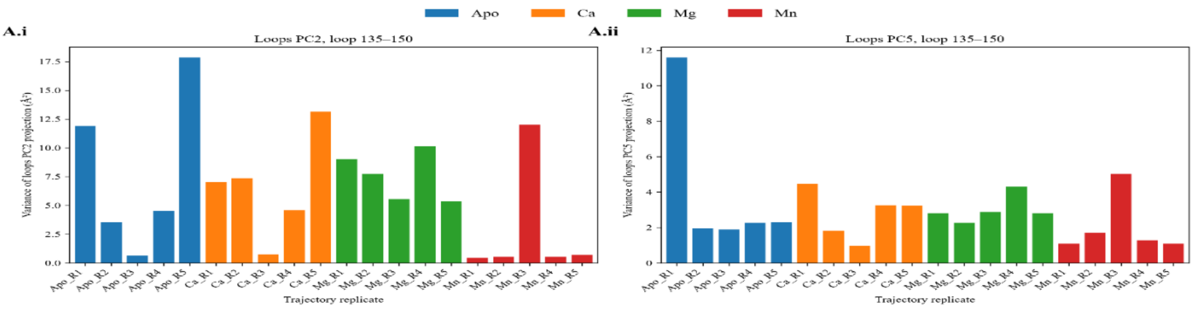


**Table 1.** Loops PC2 projection statistics of loop 135–150, all 20 replicates

| Replicate | Mean (Å) | Variance (Å²) | SD (Å) | Range (Å) |
|---|---|---|---|---|
| Apo_R1 | -1.713 | 11.893 | 3.449 | 14.835 |
| Apo_R2 | 2.978 | 3.552 | 1.885 | 10.823 |
| Apo_R3 | 3.702 | 0.634 | 0.797 | 6.688 |
| Apo_R4 | -0.560 | 4.523 | 2.127 | 11.767 |
| Apo_R5 | 0.459 | 17.863 | 4.227 | 14.462 |
| Ca_R1 | -2.507 | 7.029 | 2.651 | 13.979 |
| Ca_R2 | 2.252 | 7.367 | 2.714 | 13.480 |
| Ca_R3 | 2.954 | 0.759 | 0.871 | 6.348 |
| Ca_R4 | -3.591 | 4.598 | 2.144 | 16.029 |
| Ca_R5 | -2.395 | 13.155 | 3.627 | 17.004 |
| Mg_R1 | -2.997 | 9.025 | 3.004 | 14.914 |
| Mg_R2 | -0.641 | 7.728 | 2.780 | 13.655 |
| Mg_R3 | -3.138 | 5.562 | 2.358 | 12.646 |
| Mg_R4 | -6.362 | 10.142 | 3.185 | 19.133 |
| Mg_R5 | -1.680 | 5.332 | 2.309 | 13.344 |
| Mn_R1 | 3.385 | 0.445 | 0.667 | 5.251 |
| Mn_R2 | 3.111 | 0.528 | 0.726 | 5.206 |
| Mn_R3 | 0.259 | 11.988 | 3.462 | 13.499 |
| Mn_R4 | 3.388 | 0.525 | 0.725 | 6.776 |
| Mn_R5 | 3.093 | 0.701 | 0.838 | 6.342 |

**Table 3.** Loops PC2: mean over R1–R5 of each system

| System | Mean variance, R1–R5 (Å²) | Mean SD, R1–R5 (Å) |
|---|---|---|
| Apo | 7.693 | 2.497 |
| Ca | 6.582 | 2.402 |
| Mg | 7.558 | 2.727 |
| Mn | 2.838 | 1.284 |

**Table 2.** Loops PC5 projection statistics of loop 135–150, all 20 replicates

| Replicate | Mean (Å) | Variance (Å²) | SD (Å) | Range (Å) |
|---|---|---|---|---|
| Apo_R1 | -0.533 | 11.601 | 3.406 | 17.288 |
| Apo_R2 | -0.517 | 1.953 | 1.398 | 10.649 |
| Apo_R3 | -1.174 | 1.879 | 1.371 | 9.839 |
| Apo_R4 | 1.811 | 2.273 | 1.508 | 9.674 |
| Apo_R5 | 0.031 | 2.307 | 1.519 | 12.885 |
| Ca_R1 | -2.280 | 4.471 | 2.114 | 15.572 |
| Ca_R2 | -0.443 | 1.814 | 1.347 | 10.192 |
| Ca_R3 | -0.206 | 0.972 | 0.986 | 7.657 |
| Ca_R4 | 1.245 | 3.249 | 1.803 | 13.346 |
| Ca_R5 | 0.388 | 3.232 | 1.798 | 14.507 |
| Mg_R1 | 1.582 | 2.822 | 1.680 | 12.350 |
| Mg_R2 | 4.146 | 2.273 | 1.508 | 10.668 |
| Mg_R3 | 1.559 | 2.863 | 1.692 | 10.666 |
| Mg_R4 | 0.480 | 4.308 | 2.076 | 14.740 |
| Mg_R5 | 2.635 | 2.822 | 1.680 | 12.781 |
| Mn_R1 | -0.880 | 1.087 | 1.042 | 7.919 |
| Mn_R2 | -1.226 | 1.687 | 1.299 | 8.912 |
| Mn_R3 | -4.433 | 5.018 | 2.240 | 13.891 |
| Mn_R4 | -1.766 | 1.285 | 1.133 | 10.031 |
| Mn_R5 | -0.419 | 1.096 | 1.047 | 7.900 |

**Table 4.** Loops PC5: mean over R1–R5 of each system

| System | Mean variance, R1–R5 (Å²) | Mean SD, R1–R5 (Å) |
|---|---|---|
| Apo | 4.003 | 1.840 |
| Ca | 2.748 | 1.610 |
| Mg | 3.018 | 1.727 |
| Mn | 2.035 | 1.352 |

**Figure S10:** Histogram plots of variance of movement of loop 135-150 along PC2 and PC5 eigenmode respectively. (**Table 1 – 2)** Statistics of movement of loop 135-150 along PC2 and PC5 eigenmode respectively.

### Supplementary figure S11:

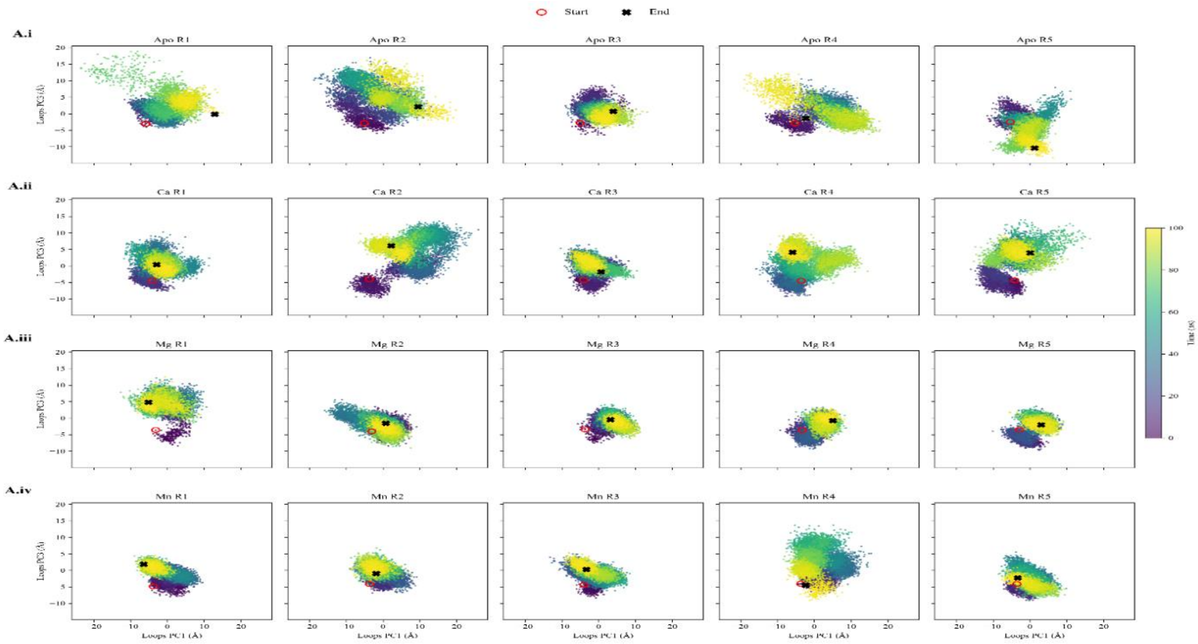

**Figure S11: (A.i-A.iv)** Projections of the trajectories of 5 runs of each state of loop 199-223 backbone (C-α atoms) onto the PC1 and PC3 eigenvector components of loop-specific combined Cartesian PCA.

**Supplementary figure S12:**

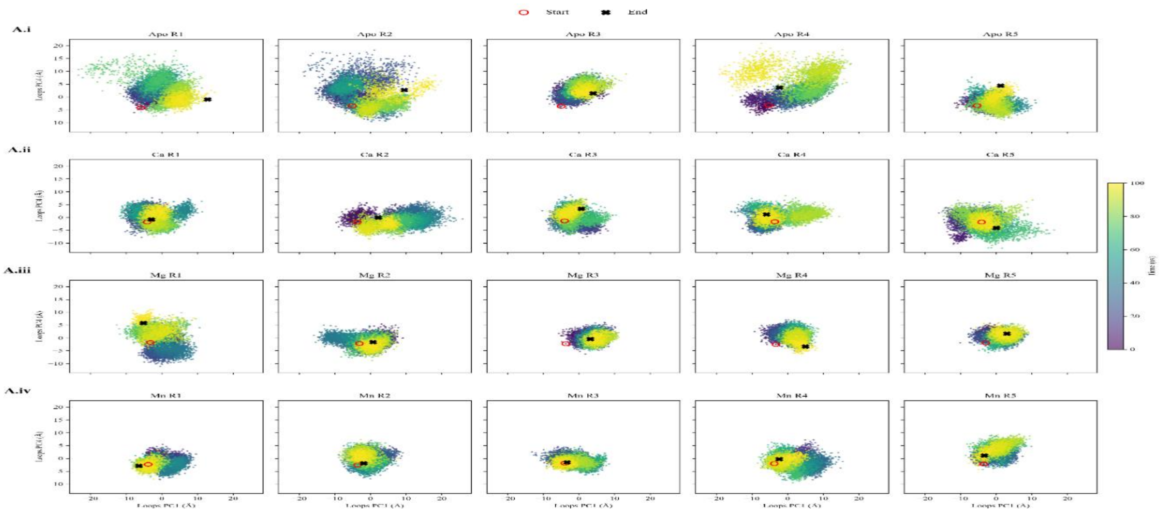


**Figure S12: (A.i-A.iv)** Projections of the trajectories of 5 runs of each state of loop199-223 backbone (C-α atoms) onto the PC1 and PC4 eigenvector components of loop-specific combined Cartesian PCA.

**Supplementary figure S13:**

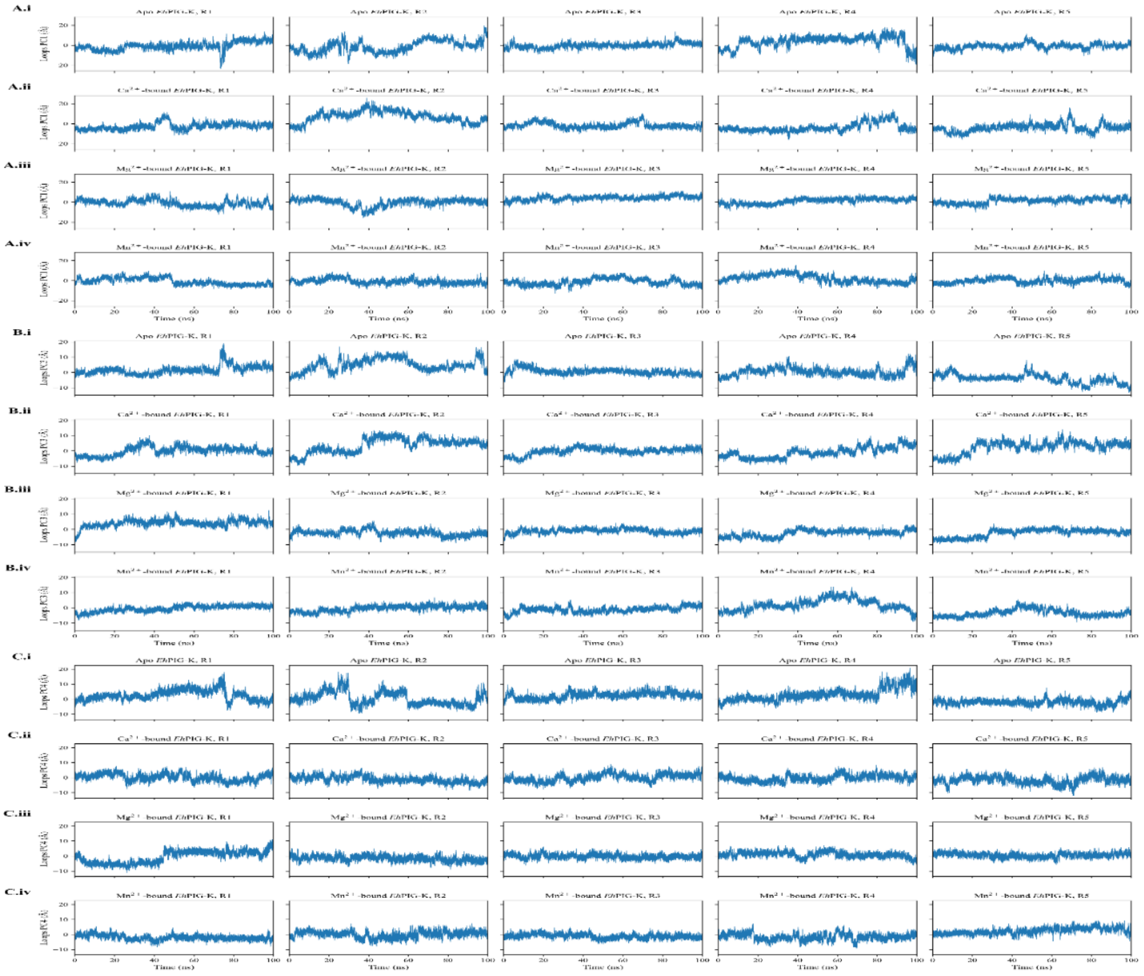

**Figure S13: (A.i-A.iv)** Time-resolved projections of loop 199-223 backbone trajectories onto PC1 eigenvector components. (**B.i-B.iv)** Time-resolved projections of loop 199-223 backbone trajectories onto PC3 eigenvector components. **(C.i-C.iv)** Time-resolved projections of loop 199-223 backbone trajectories onto PC4 eigenvector components.

### Supplementary figure S14:

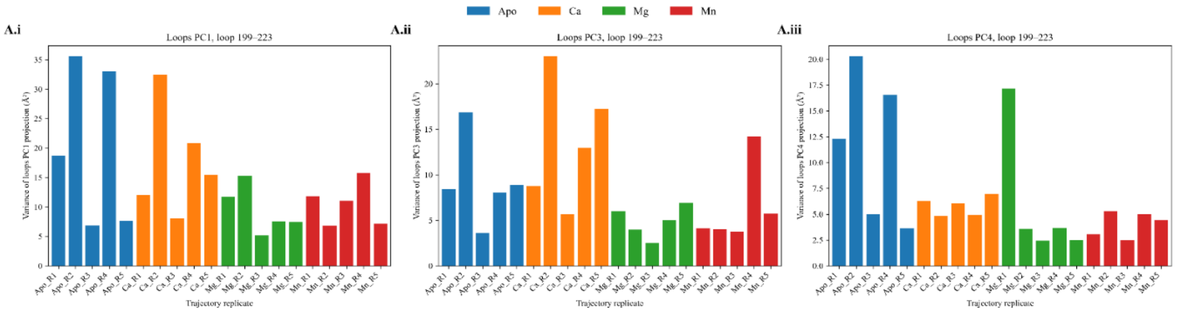


**Table 1.** Loops PC1 projection statistics of loop 199–223, all 20 replicates

| Replicate | Mean (Å) | Variance (Å²) | SD (Å) | Range (Å) |
|---|---|---|---|---|
| Apo_R1 | -0.794 | 18.713 | 4.326 | 38.300 |
| Apo_R2 | -1.186 | 35.634 | 5.969 | 38.459 |
| Apo_R3 | -0.227 | 6.890 | 2.625 | 22.174 |
| Apo_R4 | 3.604 | 33.049 | 5.749 | 36.876 |
| Apo_R5 | -0.966 | 7.602 | 2.757 | 22.697 |
| Ca_R1 | -2.592 | 12.015 | 3.466 | 23.081 |
| Ca_R2 | 7.133 | 32.442 | 5.696 | 35.455 |
| Ca_R3 | -1.690 | 8.052 | 2.838 | 20.695 |
| Ca_R4 | -3.737 | 20.832 | 4.564 | 30.303 |
| Ca_R5 | -4.015 | 15.448 | 3.930 | 32.348 |
| Mg_R1 | -0.949 | 11.708 | 3.422 | 23.254 |
| Mg_R2 | -0.383 | 15.284 | 3.909 | 24.939 |
| Mg_R3 | 4.107 | 5.157 | 2.271 | 16.823 |
| Mg_R4 | 1.137 | 7.539 | 2.746 | 17.915 |
| Mg_R5 | 1.387 | 7.420 | 2.724 | 19.226 |
| Mn_R1 | -0.730 | 11.807 | 3.436 | 18.523 |
| Mn_R2 | -0.863 | 6.800 | 2.608 | 18.048 |
| Mn_R3 | -1.279 | 11.037 | 3.322 | 21.647 |
| Mn_R4 | 2.076 | 15.771 | 3.971 | 23.816 |
| Mn_R5 | -0.037 | 7.196 | 2.682 | 18.411 |

**Table 2.** Loops PC3 projection statistics of loop 199–223, all 20 replicates

| Replicate | Mean (Å) | Variance (Å²) | SD (Å) | Range (Å) |
|---|---|---|---|---|
| Apo_R1 | 1.578 | 8.405 | 2.899 | 23.235 |
| Apo_R2 | 5.056 | 16.864 | 4.107 | 22.859 |
| Apo_R3 | 0.609 | 3.607 | 1.899 | 15.024 |
| Apo_R4 | 0.347 | 8.050 | 2.837 | 18.868 |
| Apo_R5 | -4.121 | 8.898 | 2.983 | 21.301 |
| Ca_R1 | -0.095 | 8.751 | 2.958 | 17.838 |
| Ca_R2 | 3.703 | 23.040 | 4.800 | 23.192 |
| Ca_R3 | -0.204 | 5.635 | 2.374 | 15.441 |
| Ca_R4 | -1.032 | 12.955 | 3.599 | 19.196 |
| Ca_R5 | 2.442 | 17.266 | 4.155 | 23.719 |
| Mg_R1 | 3.998 | 6.022 | 2.454 | 20.912 |
| Mg_R2 | -2.307 | 3.987 | 1.997 | 14.668 |
| Mg_R3 | -1.139 | 2.497 | 1.580 | 11.686 |
| Mg_R4 | -2.595 | 5.002 | 2.237 | 12.653 |
| Mg_R5 | -2.529 | 6.920 | 2.631 | 13.248 |
| Mn_R1 | -0.526 | 4.130 | 2.032 | 12.505 |
| Mn_R2 | -0.429 | 4.040 | 2.010 | 13.581 |
| Mn_R3 | -1.216 | 3.778 | 1.944 | 13.639 |
| Mn_R4 | 1.945 | 14.218 | 3.771 | 22.855 |
| Mn_R5 | -3.485 | 5.737 | 2.395 | 14.730 |

**Table 3.** Loops PC4 projection statistics of loop 199–223, all 20 replicates

| Replicate | Mean (Å) | Variance (Å²) | SD (Å) | Range (Å) |
|---|---|---|---|---|
| Apo_R1 | 2.715 | 12.322 | 3.510 | 25.482 |
| Apo_R2 | 0.423 | 20.318 | 4.508 | 27.813 |
| Apo_R3 | 2.462 | 4.981 | 2.232 | 15.283 |
| Apo_R4 | 3.004 | 16.564 | 4.070 | 27.971 |
| Apo_R5 | -2.036 | 3.630 | 1.905 | 16.641 |
| Ca_R1 | 0.150 | 6.275 | 2.505 | 15.413 |
| Ca_R2 | -1.210 | 4.819 | 2.195 | 14.737 |
| Ca_R3 | -0.060 | 6.051 | 2.460 | 16.544 |
| Ca_R4 | -0.398 | 4.941 | 2.223 | 15.434 |
| Ca_R5 | -1.758 | 6.979 | 2.642 | 18.758 |
| Mg_R1 | -0.574 | 17.146 | 4.141 | 22.099 |
| Mg_R2 | -1.623 | 3.568 | 1.889 | 14.139 |
| Mg_R3 | -0.206 | 2.434 | 1.560 | 12.379 |
| Mg_R4 | 0.947 | 3.666 | 1.915 | 13.149 |
| Mg_R5 | 0.491 | 2.526 | 1.589 | 11.573 |
| Mn_R1 | -1.998 | 3.068 | 1.752 | 13.749 |
| Mn_R2 | 0.012 | 5.285 | 2.299 | 15.314 |
| Mn_R3 | -1.164 | 2.493 | 1.579 | 11.108 |
| Mn_R4 | -1.575 | 5.007 | 2.238 | 16.720 |
| Mn_R5 | 2.396 | 4.433 | 2.106 | 13.882 |

**Table 4.** Loops PC1: mean over R1–R5 of each system

| System | Mean variance, R1–R5 (Å²) | Mean SD, R1–R5 (Å) |
|---|---|---|
| Apo | 20.378 | 4.285 |
| Ca | 17.758 | 4.099 |
| Mg | 9.422 | 3.014 |
| Mn | 10.522 | 3.204 |

**Table 5.** Loops PC3: mean over R1–R5 of each system

| System | Mean variance, R1–R5 (Å²) | Mean SD, R1–R5 (Å) |
|---|---|---|
| Apo | 9.165 | 2.945 |
| Ca | 13.529 | 3.577 |
| Mg | 4.886 | 2.180 |
| Mn | 6.381 | 2.430 |

**Table 6.** Loops PC4: mean over R1–R5 of each system

| System | Mean variance, R1–R5 (Å²) | Mean SD, R1–R5 (Å) |
|---|---|---|
| Apo | 11.563 | 3.245 |
| Ca | 5.813 | 2.405 |
| Mg | 5.868 | 2.219 |
| Mn | 4.057 | 1.994 |

**Figure S14: (A.i-A.iii)** Histogram plots of variance of movement of loop 199-223 along PC1, PC3 and PC4 eigenmodes respectively. (**Table 1 – 3)** Statistics of movement of loop 135-150 along PC1, PC3 and PC4 eigenmodes respectively.

**Supplementary figure S15:**

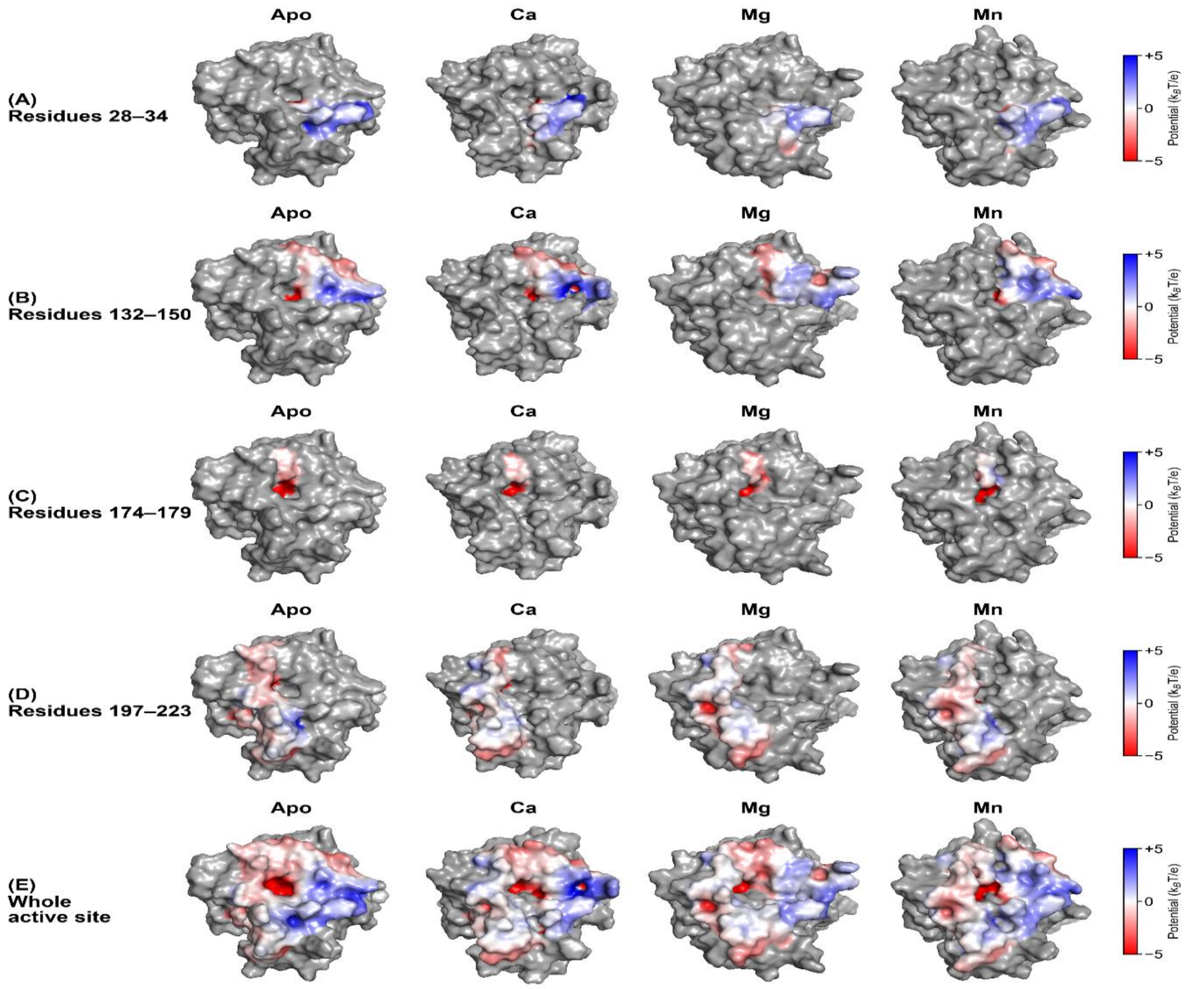


**Figure S15: (A-D)** Electrostatic contributions of regions 28-34, 132-150, 174-179 and 199-223 in consecutive sequence to the formation of the active site across all four states. **(E)** Electrostatic mapping of the active site across all four states.
